# Identification of the client-binding site on the Golgi membrane protein adaptor Vps74

**DOI:** 10.1101/2025.02.10.637573

**Authors:** Agnieszka Lesniak, Ziyun Ye, David K. Banfield

## Abstract

Vps74 and its mammalian counterpart GOLPH3 are COPI associated protein sorting adaptors that function to maintain the cisternal distributions of certain Golgi integral membrane protein clients. The GOLPH3 adaptors accommodate a diversity of clients by binding to their short cytoplasmically exposed N-termini. Here, we identify the client-binding site on yeast GOLPH3 (Vps74) which maps to two evolutionarily conserved unstructured regions on the membrane-facing surface of the protein. The client-binding site includes residues previously shown to mediate binding of GOLPH3s to PI4P, as well as the membrane-binding β-hairpin. Clients have varying requirements for the binding interface, and our study thus reconciles how a diversity of client N-termini can be accommodated by Vps74. We establish that the binding site for the PI4P phosphatase Sac1 and the COPI-coatomer associated GTPase Arf1 overlaps with and obscures client access, suggesting a regulatory role for these proteins in opposition to the adaptor. Furthermore, we identify an additional mode for the recruitment of the adaptor from the cytoplasm to Golgi membranes whereby Vps74 binds directly to its client N-termini.

## INTRODUCTION

The Golgi is the protein sorting hub of the cell, and the functional and compositional integrity of this organelle relies on its capacity to discriminate resident from non-resident proteins (Banfield, 2011; Tu and Banfield, 2010). Golgi resident proteins are typically integral membrane proteins with roles in glycosylation of proteins and lipids, ion transport, proteolytic processing, and the synthesis and processing of lipids. In contrast to other intra-cellular organelles the Golgi is comprised of several sub-compartments termed cisterna and newly synthesized proteins are transported vectorially from *cis*, medial to *trans* whereupon they are successively and sequentially modified. The enzymes that modify nascent proteins are enriched in the cisterna in which they function, and thus in addition to discriminating between resident and non-resident proteins the Golgi also needs to maintain the correct cisternal distributions of its native proteins (Banfield, 2011; Tu and Banfield, 2010).

A prevailing model for the retention of Golgi proteins is the cisternal maturation model which stipulates that cisternal identity is maintained through iterative cycles of capture and retrieval of resident proteins from *trans* to *cis* cisternae mediated in COPI-coatomer coated vesicles (Glick and Nakano, 2009; Glick and Luini; 2011; Pantazopoulou and Glick; 2019). The key question in understanding the maintenance of the compartmental protein composition of the Golgi has been to address how resident proteins are recognized, selected and sorted into COPI vesicles. In addition to informing fundamental cell biology, defects in Golgi function are associated with human diseases including congenital defects in glycosylation, neurodegenerative diseases as well as a variety of cancers (Li et al., 2019).

To date, several Golgi protein sorting adaptors have been identified and they include GOLPH3 (Vps74 in yeast cells) (Rizzo et al., 2021; Schmitz et al., 2008; Tu et al., 2008 and Welch et al., 2021), Erd1 (Sardana et al., 2021), FAM114A2 (Welch et al., 2024) and most recently LYSET (Brauer et al., 2024). In the case of the GOLPH3s, the adaptor binds to comparatively short, cytoplasmically exposed N-termini of certain Golgi membrane proteins. The N-termini of GOLPH3’s clients vary somewhat in their length and amino acid sequence but in general appear to be comprised of a cluster of hydrophobic residues, or often positively charged amino acids such as lysine and arginine (Rizzo et al., 2021; Tu et al., 2008; Welch et al., 2021). Like GOLPH3s, FAM114A2 proteins appear to recognize a cluster of basic amino acids as well (Welch et al., 2024), whereas the features of Golgi residents recognized by Erd1 are presently unknown. LYSET is an atypical GOLPH3 client whose Golgi retention is critical for the sorting of some proteins to the lysosome (Brauer et al., 2024). GOLPH3s bind directly to coatomer providing a mechanistic link between client-binding and incorporation into COPI vesicles (Eckert et al., 2014; Tu et al., 2012; Welch et al., 2021).

GOLPH3 is an oncogene (Scott et al., 2009). Amplification of GOLPH3 is observed in several cancers where it is associated with poor prognosis, revealing an important role for this Golgi protein in cancer progression (Rizzo et al., 2019; Sechi et al., 2020). It was initially suggested that the oncogenic properties of GOLPH3 were related to enhanced activation of mTOR (Scott et al., 2009; Sechi et al., 2015). However, more recently, the oncogenic role of GOLPH3 has been shown to involve its requirement for the retention of certain Golgi membrane proteins (Rizzo et al., 2021).

The crystal structures of yeast and human GOLPH3s have been determined, and they share a striking degree of similarity in their folded domain (Schmitz et al., 2008; Wood et al., 2009). From these structures it was surmised that GOLPH3s bind to PI4P, a phosphoinositide that is enriched in Golgi *trans* cisternae and in the *trans* Golgi network (TGN). Amino acid substitutions at the presumptive PI4P binding site resulted in loss of Golgi membrane binding and concomitant mislocalization of Vps74 clients from the Golgi to the vacuole. Further interrogation of features of the folded domain of Vps74 / GOLPH3s revealed a role for the conserved β hairpin in membrane-binding. Thus, PI4P-binding and the hydrophobic tip of the protein’s β hairpin are likely to account for the GOLPH3s Golgi membrane localization. Although it is not known whether either determinant will suffice or if PI4P and the β hairpin are both required for membrane recruitment.

Further structural and biochemical studies established that GOLPH3s bind to Sac1 (a PI4P phosphatase) and that GOLPH3s are Sac1 effector proteins (Cai et al., 2014; Wood et al., 2012). However, it remains to be established to what extent PI4P binding participates in adaptor - client-binding. The N-terminal region of the GOLPH3s varies in length from species to species and is likely to be intrinsically unstructured, nevertheless this region contains an evolutionarily conserved cluster of arginine residues that are critical for the adaptor’s binding to coatomer (Tu et al., 2012). Biochemical studies have shown that Vps74 binds to the coatomer-associated small GTPase Arf1 (Tu et al., 2012). The association of Arf1 and Vps74 is mediated by the folded domain of Vps74 and requires that Arf1 be bound to GTP. The functional relevance of this interaction has not been investigated, but the necessity for Arf1-GTP suggests that the interaction with Vps74 occurs on Golgi membranes.

Importantly, it is not presently known where on GOLPH3s clients bind. This information is critical for extending our understanding of the molecular mechanisms by which membrane proteins are retained in the Golgi and will also inform therapeutic intervention strategies for GOLPH3s in cancer cells.

Herein we report identification of the client-binding site on the yeast GOLPH3 adaptor Vps74. Client-binding is mediated through two evolutionarily conserved membrane opposing loops, the PI4P binding site and the β hairpin suggesting that the mode of client-adaptor binding is likely to be conserved amongst GOLPH3s. Moreover, we establish that client over-expression is sufficient to recruit Vps74 to the Golgi. We further show that the binding site for clients on Vps74 overlaps with those for Sac1 and Arf1-GTP, findings that have important implications for the roles of Sac1 and Arf1-GTP in GOLPH3-mediated retention of Golgi membrane proteins.

## MATERIALS AND METHODS

### MATERIALS

### METHODS

#### Bioinformatics

The structures of Vps74 individually bound to nine of its client N-termini (Figure EV3A) were generated with Alphafold2 multimer powered by the COSMIC2 platform (https://cosmic-cryoem.org). Three models from each prediction were analyzed using the software ChimeraX (https://www.rbvi.ucsf.edu/chimerax). The intermolecular interactions between Vps74 and client N-termini were determined with the Contacts tool (VDW overlap≥-0.40Å) and the interaction frequency of every residue in Vps74 was recorded.

The predicted structure of Vps74 bound to Vps74InNb#4 was generated with Alphafold3 (https://alphafoldserver.com/about).

#### Yeast cell serial dilutions

Yeast cells were grown to early-mid log phase (OD_660_ ∼0.5 - 0.8) and ∼10^7^ cells were collected by centrifugation. Cell pellets were resuspended in ddH_2_O for 10-fold serial dilutions and 1/10 of the cell suspension was spotted on agar plates (∼10^6^ cells in the first spot). Images of plates were taken after 48 - 72 h of incubation at the indicated temperatures. For serial dilutions conducted on calcofluor white (CFW) plates, 50 μg/mL CFW was added to the YEPD medium. For serial dilutions conducted on 5-fluoroorotic acid (5-FOA) plates, 1 mg/mL 5-FOA was added to the synthetic defined (SD) medium.

#### Preparation of yeast whole cell extracts for immunoblotting of Vps74 at 25°C and 37°C

For Figure 1D, the indicated yeast strains were grown in YEPD at 25°C to an OD_660_ ∼0.5 after which 10^7^ cells were harvested by centrifugation and resuspended in 15% v/v trichloroacetic acid (TCA). The remaining culture was diluted in pre-warmed YEPD (37°C) to ∼0.2×10^7^ cells/mL and thereafter were incubated with shaking at 37°C for 3 hours. Cells were collected by centrifugation and resuspended in 15% v/v TCA. The cell / TCA suspensions were incubated overnight at -20°C, protein precipitates were collected by centrifugation, and protein pellets were washed once with cold acetone (-20°C). Pellets were air dried and solubilized in 100 μl SDS-PAGE sample buffer containing 2.5% SDS, 50 mM NaOH and 5 mM DTT, and thereafter heated to 95°C for 10 min. Proteins were resolved by SDS-PAGE and immunoblotted with an anti-Vps74 antibody (Table 5).

**Figure 1.**
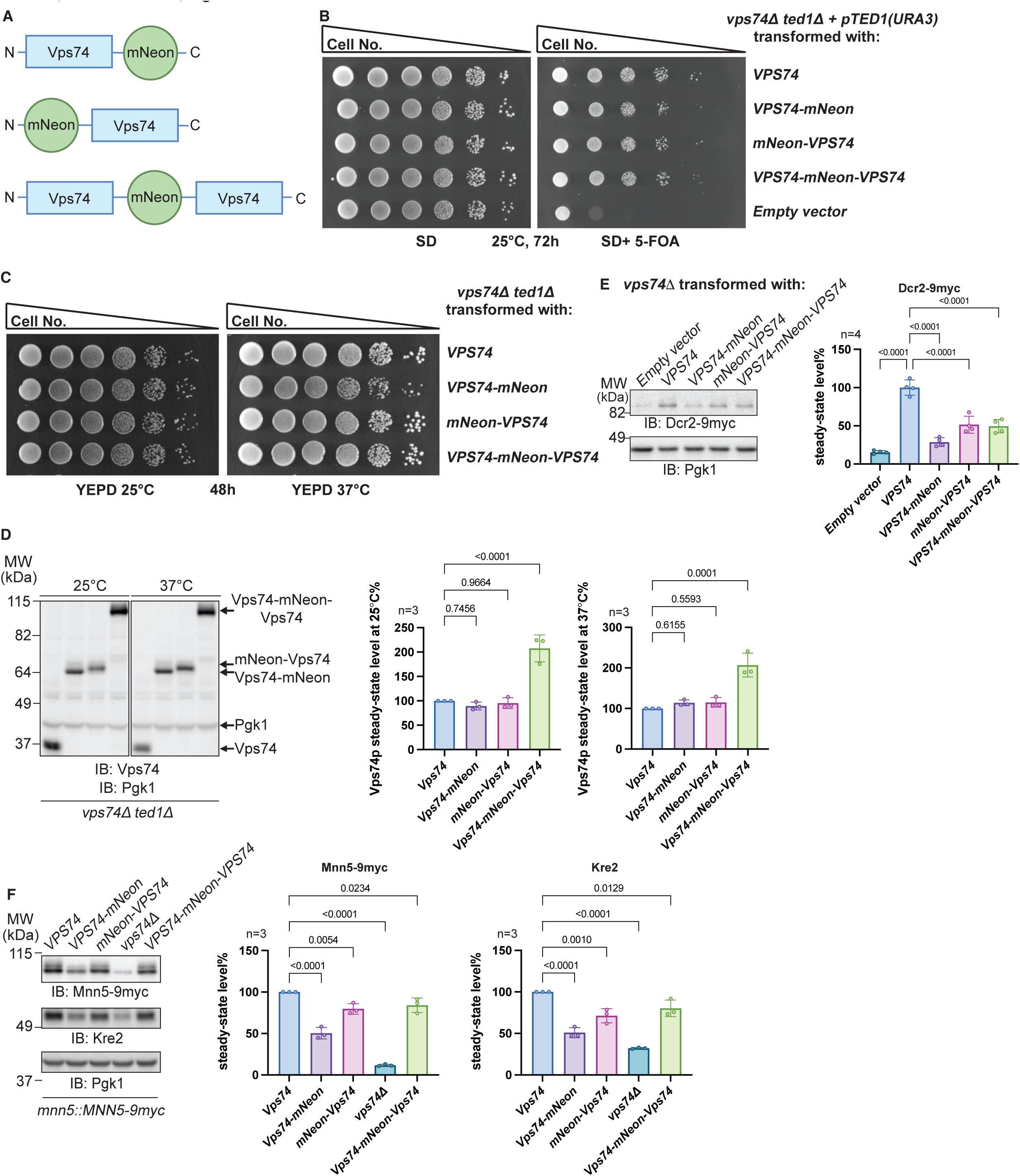
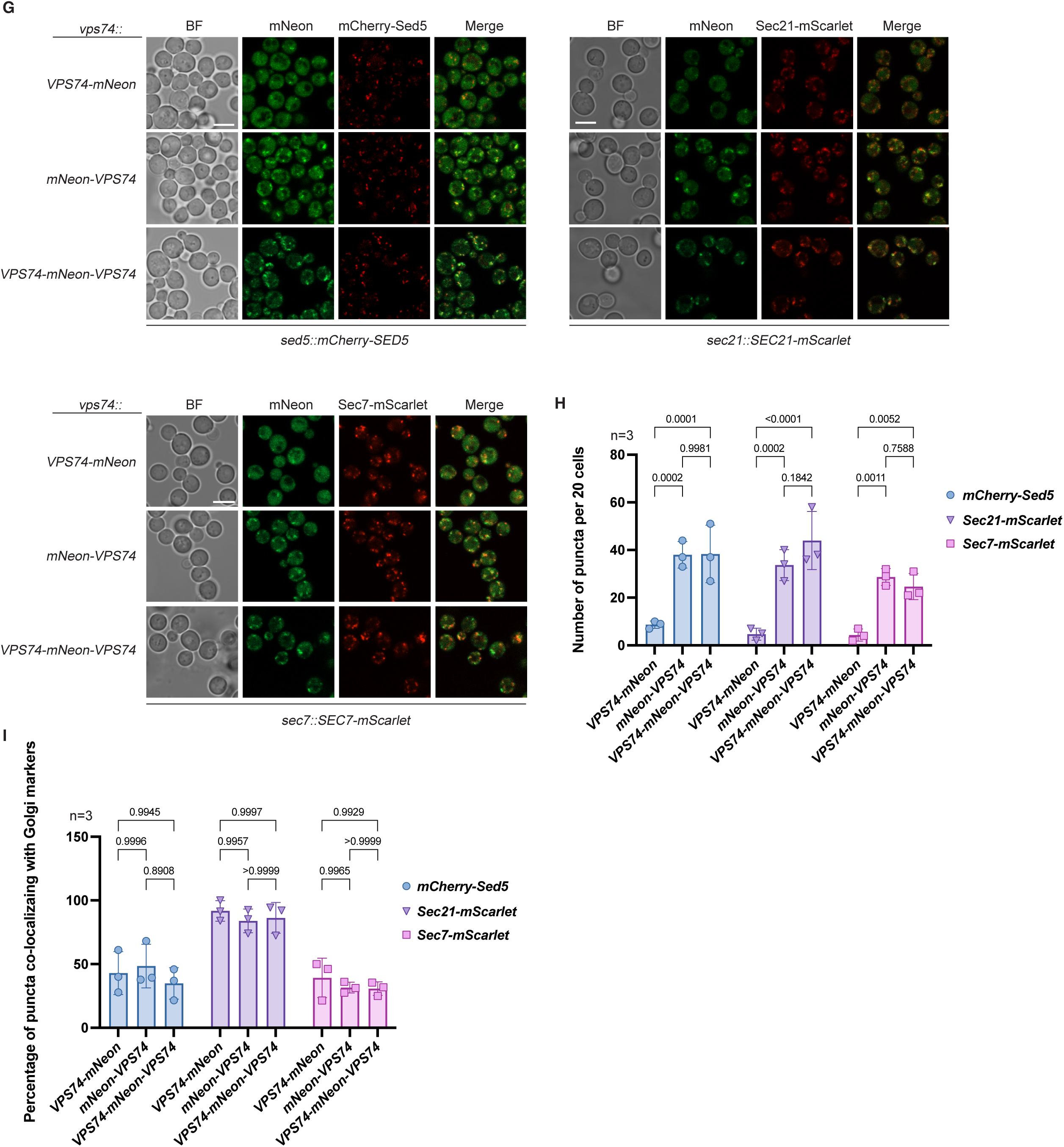
Vps74 / mNeon fusion proteins do not differ in their Golgi cisternae distributions. (A) Schematic representations of the various Vps74 / mNeon fusion proteins used in this study. (B) *VPS74*-mNeon, mNeon-*VPS74* and *VPS74*-mNeon-*VPS74* can support the growth of the synthetic deletion mutant *vps7411 ted111*. 10-fold serial dilutions of the indicated yeast strains were spotted onto plates and thereafter incubated as indicated. (C) *vps7411 ted111* cells expressing the various Vps74 / mNeon fusion proteins are not temperature-sensitive for growth. 10-fold serial dilutions of the indicated yeast strains were spotted onto plates and thereafter incubated as indicated. (D) Immunoblot and quantification of the steady-state levels of the various Vps74 / mNeon fusion proteins in *vps7411 ted111* cells. Pgk1 serves as a gel load control. (E) Immunoblots and quantification of steady-state levels of Dcr2 in *vps7411* cells expressing the various Vps74 / mNeon fusion proteins. Pgk1 serves as a gel load control. (F) Immunoblots and quantification of the steady-state levels of Mnn5 and Kre2 in cells expressing the various Vps74 / mNeon fusion proteins. Pgk1 serves as a gel load control. (G) Co-localization of the various Vps74 / mNeon proteins in cells expressing mCherry-Sed5 (*cis*), Sec21-mScarlet (*cis*, medial and *trans*) or Sec7-mScarlet (*trans)* Golgi residents. Scale bar 5 μm. (H) Quantification of the number of mNeon puncta from experiments depicted in (G). (I) Quantification of the co-localization data for the various Vps74 / mNeon proteins with *cis*, medial and *trans* Golgi markers from experiments depicted in (G).

#### Measurement of steady-state levels of Golgi proteins in whole cell extracts by immunoblotting

Two different methods were employed to measure the steady-state levels of the enzymes. For Figures 1F, 4C, Figure EV1A and Figure EV2, yeast cells were grown to the early-mid log phase (OD_660_=0.5-0.8) and 10^7^ cells were collected by centrifugation. Cells were resuspended with 200 μl 100 mM NaOH and incubated for 10 min. Excess NaOH was removed by centrifugation, and cells were resuspended with 100 μl SDS sample buffer supplemented with 1x protease inhibitor cocktail, 1 mM Pefabloc SC and 5 mM DTT. Samples were then boiled for 10 min at 95°C. For the remaining Figures, the alkaline lysis step was omitted. Instead, cell pellets were directly resuspended in 100 μl SDS sample buffer supplemented with 1x protease inhibitor cocktail, 1 mM Pefabloc SC and 5 mM DTT. ∼ 30 μl acid-washed glass beads were added to cell suspensions and yeast cells were lysed by vortexing for 5 min at room temperature. Following vortexing, samples were heated at 95°C for 10 min and, and thereafter proteins were resolved by SDS-PAGE.

For the immunoblots of Dcr2, Kre2, Mnn5, Mnn4 and Och1, lysates were treated with Endoglycosidase H (Endo H). ∼20 μl of lysates in SDS sample buffer were supplemented with 3.8 μl 0.5 M potassium acetate (pH 5.6) and 10 NEB units of Endo H and incubated for 2 h at 37°C. Prior to SDS-PAGE, samples were incubated at 95°C for 5 min.

#### SDS-PAGE and immunoblotting

∼10 μl of prepared samples were loaded on 8 cm x 9cm Tris-glycine polyacrylamide gels and proteins were resolved in Tris-glycine running buffer for 1h at 200 V. For Coomassie Blue staining, gels were incubated in the staining solution (0.025% w/v Coomassie Blue R-250, 40% v/v methanol, 7% v/v acetic acid) with gentle mixing for 1 h. Gels were destained with 10% v/v acetic acid, 50% v/v methanol before imaging using a ChemiDoc Imaging System. For immunoblotting, proteins were transferred to 0.45 μm cellulose membranes, stained with Ponceau S solution (0.1% w/v Ponceau S, 5% v/v acetic acid) and images acquired with a ChemiDoc Imaging System. The stain was removed by washing the membrane in PBST (137 mM NaCl, 2.7 mM KCl, 10 mM Na_2_PO_4_, 1.8 mM KH_2_PO_4_, 0.1% v/v Tween 20) before immunoblotting. Prior to the addition of antibodies membranes were incubated in 5% w/v non-fat milk for 1 h at room temperature on an orbital shaker. Membranes were incubated with primary antibodies in 5% w/v milk for 1 h at room temperature on an orbital shaker. Excess primary antibodies were removed by washing with PBST for 5 min, three times. Membranes were incubated with secondary antibodies in 5% w/v milk for 1 h at room temperature. After washing with PBST for 5 min, three times, ECL substrate was added to membranes and chemiluminescent images acquired using a ChemiDoc Imaging System. For the immunoblotting of Mnn4-9myc, SuperSignal™ West Femto Maximum Sensitivity Substrate was used to enhance the weak signal.

#### Expression and purification of recombinant proteins from bacteria

For recombinant protein production *E. coli* BL21(DE3) cells were transformed with the various expression plasmids. LB media was inoculated with a single bacterial colony and cultures were incubated with shaking (200 rpm) at 37°C overnight (12 – 14 hours). Overnight cultures were diluted 1:40 with LB media and thereafter grown at 37°C and 200 rpm until the OD_600_ reached ∼0.6 - 0.8. Protein expression was induced with 0.2 mM IPTG at 25°C with shaking (200 rpm) overnight. Cells from 100 ml cultures were collected by centrifugation at 4,200 rcf for 10 min and washed once with PBS.

For GST-tagged fusion proteins, cell pellets were resuspended in 2 ml PBS + 10% glycerol supplemented with 1x protease inhibitor cocktail, 1 mM Pefabloc SC and 250 μg/mL lysozyme, and lysed by sonication on ice with a Q125 sonicator for 4 min, 2s on, 2s off at 40% amplitude. Following sonication cell debris was removed by centrifugation (16,000 rcf at 4°C). To estimate the relative amounts of GST-fusion proteins in the bacterial lysates, different amount of lysates (∼3-7 μl) were diluted in 100 μl PBS + 0.1% Triton X-100 and mixed with 20 μl Glutathione Sepharose 4B equilibrated with PBS + 0.1% Triton X-100 for 1 h at 4°C. After washing with 200 μl PBS + 0.1% Triton-X-100 three times, the beads were mixed with 30 μl SDS sample buffer supplemented with 5 mM DTT, heated at 95°C for 5 min and the amount of bound proteins was estimated by SDS-PAGE.

For (His)_6_-tagged fusion proteins, cell pellets were resuspended in 2 ml Ni-NTA lysis buffer (50 mM NaH_2_PO_4_, 200 mM NaCl, 10 mM imidazole, pH 8.0) supplemented with 1x protease inhibitor cocktail, 1 mM Pefabloc SC and 250 μg/mL lysozyme, and cells were lysed on ice by sonication. Debris and unlysed cells were removed by centrifugation at 16,000 rcf at 4°C for 10 min and supernatants were mixed with 0.5 ml Ni-NTA beads equilibrated with Ni-NTA lysis buffer. After 1 h mixing at 4°C, the beads were washed with 3 ml Ni-NTA wash buffer (50 mM NaH_2_PO_4_, 200 mM NaCl, 20 mM imidazole, pH 8.0) three times. Bound proteins were eluted with 500 μl PBS pH 7.4 + 250 mM imidazole, five times.

For Twin-Strep-tagged fusion proteins, proteins were purified on Strep-tactin Sepharose beads as described for (His)_6_-tagged proteins except that PBS + 1mM EDTA was used as the lysis and wash buffer, and PBS + 2.5 mM desthiobiotin was used to elute bound proteins.

The expression and purification of bacterially expressed N-myristoylated Arf1 was performed as previously described (Tu *et al*., 2012).

#### Isolation of anti-Vps74 nanobodies from the synthetic yeast surface-display library

The isolation of anti-Vps74 nanobodies was performed as previously described (McMahon et al., 2018) with the following modifications. N-terminally Twin-Strep-tagged mNeon and Twin-Strep-tagged Vps74 were used in the negative and positive selection steps, respectively. For the first round of selection, 1 ml Strep-tactin Sepharose bound to 1 mg of purified Twin-Strep-tagged mNeon or Twin-Strep-tagged Vps74 was added to an Econo-Column Chromatography Column (1.0 × 10 cm). Yeast cells displaying nanobodies against Strep-tactin Sepharose and Twin-Strep-mNeon were removed by passing 5 × 10^9^ yeast cells resuspended in 10 ml PBS + 0.1% BSA over 1 ml Strep-tactin Sepharose beads bound to purified Twin-Strep-tagged mNeon. The resulting flow through was then incubated with 1 ml of Strep-tactin Sepharose beads bound to Twin-Strep-tagged Vps74 for 1 h 4°C. Bound cells were eluted from the Twin-Strep-Vps74 column with 1 mM biotin. For the second-round selection, 100 μg Twin-Strep-Vps74 protein was bound to 100 μl Strep-tactin magnetic beads and mixed with 3 × 10^7^ cells generated by culturing cells obtained following the first round of enrichment. A third round of nanobody selection was conducted as per the second round except that 30 μl Strep-tactin magnetic beads loaded with Twin-Strep-Vps74 were incubated with 6 × 10^7^ cells generated by culturing cells obtained following the second round of enrichment. Yeast cells recovered from the third round of selection were spread onto the surface of -Trp plates containing 2% galactose. Cells from single colonies were incubated with 0.5 μg/μl 6His-mNeon-

Vps74 fusion protein in PBS for 1 h, at 4°C, washed with PBS once and examined by fluorescent microscopy to identify cells expressing anti-Vps74 nanobodies.

#### Screening of inhibitory nanobodies against Vps74 in *ted1Δ* cells

Total DNA was extracted from yeast cells recovered after three rounds of selection and used as template to amplify nanobody-HA coding sequences by the PCR using the primers Ubi-Nb G3 and HA stop TC R. The myc-ubiquitin sequence was amplified by the PCR using the primers TPI-myc G1 and Ubi-Nb G2. The two purified PCR products were used as templates and amplified by the PCR using TPI-myc G1 and HA stop TC R to generate the myc-ubiquitin-nanobody-HA DNA fragment with flanking complementary sequences to the *TPI1* promoter and *CYC1* terminator. Primer information can be found in Table 4.

**Table 1.** Yeast strains used in this study.

| Strain | Genotype | Source |
| --- | --- | --- |
| **Figure 1** |  |  |
| SARY8388 | *MATa his3Δ1 leu2Δ0 met15Δ0 ura3Δ0 ted1::kanMX4 vps74::loxP* transformed with *gTED1-pRS416* | This study |
| SARY8417 | *MATa his3Δ1 leu2Δ0 met15Δ0 ura3Δ0 ted1::kanMX4 vps74::loxP* transformed with *gVPS74-pRS415* | This study |
| SARY8421 | *MATa his3Δ1 leu2Δ0 met15Δ0 ura3Δ0 ted1::kanMX4 vps74::loxP* transformed with *gVPS74-mNeon-pRS415* | This study |
| SARY8753 | *MATa his3Δ1 leu2Δ0 met15Δ0 ura3Δ0 ted1::kanMX4 vps74::loxP* transformed with *gmNeon-VPS74-pRS415* | This study |
| SARY8755 | *MATa his3Δ1 leu2Δ0 met15Δ0 ura3Δ0 ted1::kanMX4 vps74::loxP* transformed with *gVPS74-mNeon-VPS74-pRS415* | This study |
| SARY8751 | *MATa his3Δ1 leu2Δ0 met15Δ0 ura3Δ0 vps74::kanMX4 mnn5::MNN5-9myc-loxP-HIS3-loxP* | This study |
| SARY8771 | *MATa his3Δ1 leu2Δ0 met15Δ0 ura3Δ0 vps74::KANMX4 mnn5::MNN5-9myc-loxP-HIS3-loxP pRS406-gVPS74* | This study |
| SARY8780 | *MATa his3Δ1 leu2Δ0 met15Δ0 ura3Δ0 vps74::KANMX4 mnn5::MNN5-9myc-loxP-HIS3-loxP pRS406-gVPS74-mNeon* | This study |
| SARY8783 | *MATa his3Δ1 leu2Δ0 met15Δ0 ura3Δ0 vps74::KANMX4 mnn5::MNN5-9myc-loxP-HIS3-loxP pRS406-gmNeon-VPS74* | This study |
| SARY8786 | *MATa his3Δ1 leu2Δ0 met15Δ0 ura3Δ0 vps74::KANMX4 mnn5::MNN5-9myc-loxP-HIS3-loxP pRS406-gVPS74-mNeon-VPS74* | This study |
| SARY8878 | *MATa his3Δ1 leu2Δ0 met15Δ0 ura3Δ0 vps74::gVPS74-mNeon sed5::gmCherry-SED5* | This study |
| SARY8881 | *MATa his3Δ1 leu2Δ0 met15Δ0 ura3Δ0 vps74::gmNeon-VPS74 sed5::gmCherry-SED5* | This study |
| SARY8882 | *MATa his3Δ1 leu2Δ0 met15Δ0 ura3Δ0 vps74::gVPS74-mNeon-VPS74 sed5::gmCherry-SED5* | This study |
| SARY8950 | *MATa his3Δ1 leu2Δ0 met15Δ0 ura3Δ0 vps74::gVPS74-mNeon sec21::SEC21-mScarlet-loxP-HIS3-loxP* | This study |
| SARY8967 | *MATa his3Δ1 leu2Δ0 met15Δ0 ura3Δ0 vps74::gmNeon-VPS74 sec21::SEC21-mScarlet-loxP-HIS3-loxP* | This study |
| SARY8969 | *MATa his3Δ1 leu2Δ0 met15Δ0 ura3Δ0 vps74::gVPS74-mNeon-VPS74 sec21::SEC21-mScarlet-loxP-HIS3-loxP* | This study |
| SARY8862 | *MATa his3Δ1 leu2Δ0 met15Δ0 ura3Δ0 vps74::gVPS74-mNeon sec7::SEC7-mScarlet-loxP-HIS3-loxP* | This study |
| SARY8864 | *MATa his3Δ1 leu2Δ0 met15Δ0 ura3Δ0 vps74::gmNeon-VPS74 sec7::SEC7-mScarlet-loxP-HIS3-loxP* | This study |
| SARY8866 | *MATa his3Δ1 leu2Δ0 met15Δ0 ura3Δ0 vps74::gVPS74-mNeon-VPS74 sec7::SEC7-mScarlet-loxP-HIS3-loxP* | This study |
| **Figure 2** |  |  |
| SARY3739 | *MATa his3Δ1 leu2Δ0 met15Δ0 ura3Δ0 vps74::kanMX4 ted1::clonNAT* transformed with *gvps74(6AAA8)-pRS416* | Lab collection |
| SARY8878 | *MATa his3Δ1 leu2Δ0 met15Δ0 ura3Δ0 vps74::gVPS74-mNeon sed5::gmCherry-SED5* | This study |
| SARY8950 | *MATa his3Δ1 leu2Δ0 met15Δ0 ura3Δ0 vps74::gVPS74-mNeon sec21::SEC21-mScarlet-loxP-HIS3-loxP* | This study |
| SARY8862 | *MATa his3Δ1 leu2Δ0 met15Δ0 ura3Δ0 vps74::gVPS74-mNeon sec7::SEC7-mScarlet-loxP-HIS3-loxP* | This study |
| SARY9118 | *MATa his3Δ1 leu2Δ0 met15Δ0 ura3Δ0 grx7::GRX7-9myc-loxP-HIS3-loxP* | This study |
| SARY9121 | *MATa his3Δ1 leu2Δ0 met15Δ0 ura3Δ0 vps74::kanMX4 grx7::GRX7-9myc-loxP-HIS3-loxP* | This study |
| **Figure 3** |  |  |
| SARY8388 | *MATa his3Δ1 leu2Δ0 met15Δ0 ura3Δ0 ted1::kanMX4 vps74::loxP* transformed with *gTED1-pRS416* | This study |
| SARY8759 | *MATa his3Δ1 leu2Δ0 met15Δ0 ura3Δ0 ted1::kanMX4 vps74::loxP* transformed with *gVPS74-pRS415* | This study |
| SARY8761 | *MATa his3Δ1 leu2Δ0 met15Δ0 ura3Δ0 ted1::kanMX4 vps74::loxP* transformed with *gvps74(W88A)-pRS415* | This study |
| SARY8763 | *MATa his3Δ1 leu2Δ0 met15Δ0 ura3Δ0 ted1::kanMX4 vps74::loxP* transformed with *gvps74(K178A, R181A)-pRS415* | This study |
| SARY8765 | *MATa his3Δ1 leu2Δ0 met15Δ0 ura3Δ0 ted1::kanMX4 vps74::loxP* transformed with *gvps74(Δ197-208)-pRS415* | This study |
| SARY8627 | *MATa his3Δ1 leu2Δ0 met15Δ0 ura3Δ0 vps74::gVPS74-mNeon* | This study |
| SARY9011 | *MATa his3Δ1 leu2Δ0 met15Δ0 ura3Δ0 vps74::gvps74(W88A)-mNeon* | This study |
| SARY9019 | *MATa his3Δ1 leu2Δ0 met15Δ0 ura3Δ0 vps74::gvps74(R97A)-mNeon* | This study |
| SARY8598 | *MATa his3Δ1 leu2Δ0 met15Δ0 ura3Δ0 vps74::KANMX4 PRS406-gvps74(K178A, R181A)-mNeon* | This study |
| SARY9024 | *MATa his3Δ1 leu2Δ0 met15Δ0 ura3Δ0 vps74::gvps74(Δ197-208)-mNeon* | This study |
| SARY8982 | *MATa his3Δ1 leu2Δ0 met15Δ0 ura3Δ0 vps74::kanMX4 mnn2::MNN2-9myc-loxP mnn4::MNN4-9myc-loxP* transformed with *gGRX7-4HA-pRS416* | This study |
| SARY8751 | *MATa his3Δ1 leu2Δ0 met15Δ0 ura3Δ0 vps74::kanMX4 mnn5::MNN5-9myc-loxP-HIS3-loxP* | This study |
| SARY9104 | *MATa his3Δ1 leu2Δ0 met15Δ0 ura3Δ0 vps74::kanMX4 dcr2::DCR2-9myc-loxP-HIS3-loxP* | This study |
| SARY9121 | *MATa his3Δ1 leu2Δ0 met15Δ0 ura3Δ0 vps74::kanMX4 grx7::GRX7-9myc-loxP-HIS3-loxP* | This study |
| **Figure 4** |  |  |
| SARY9030 | *MATa his3Δ1 leu2Δ0 met15Δ0 ura3Δ0 vps74::kanMX4 mnn5::MNN5-9myc-loxP-HIS3-loxP pRS406-gVPS74-mNeon-VPS74* | This study |
| SARY9032 | *MATa his3Δ1 leu2Δ0 met15Δ0 ura3Δ0 vps74::kanMX4 mnn5::MNN5-9myc-loxP-HIS3-loxP pRS406-gVPS74-mNeon-PH-VPS74* | This study |
| SARY9034 | *MATa his3Δ1 leu2Δ0 met15Δ0 ura3Δ0 vps74::kanMX4 mnn5::MNN5-9myc-loxP-HIS3-loxP pRS406-gvps74(W88A)-mNeon-vps74(W88A)* | This study |
| SARY9036 | *MATa his3Δ1 leu2Δ0 met15Δ0 ura3Δ0 vps74::kanMX4 mnn5::MNN5-9myc-loxP-HIS3-loxP pRS406-gvps74(W88A)-mNeon-PH-vps74(W88A)* | This study |
| SARY9038 | *MATa his3Δ1 leu2Δ0 met15Δ0 ura3Δ0 vps74::kanMX4 mnn5::MNN5-9myc-loxP-HIS3-loxP pRS406-gvps74(R97A)-mNeon-vps74(R97A)* | This study |
| SARY9040 | *MATa his3Δ1 leu2Δ0 met15Δ0 ura3Δ0 vps74::kanMX4 mnn5::MNN5-9myc-loxP-HIS3-loxP pRS406-gvps74(R97A)-mNeon-PH-vps74(R97A)* | This study |
| SARY9042 | *MATa his3Δ1 leu2Δ0 met15Δ0 ura3Δ0 vps74::kanMX4 mnn5::MNN5-9myc-loxP-HIS3-loxP pRS406-gvps74(K178A, R181A)-mNeon-vps74(K178A, R181A)* | This study |
| SARY9044 | *MATa his3Δ1 leu2Δ0 met15Δ0 ura3Δ0 vps74::kanMX4 mnn5::MNN5-9myc-loxP-HIS3-loxP pRS406-gvps74(K178A, R181A)-mNeon-PH-vps74(K178A, R181A)* | This study |
| SARY9046 | *MATa his3Δ1 leu2Δ0 met15Δ0 ura3Δ0 vps74::kanMX4 mnn5::MNN5-9myc-loxP-HIS3-loxP pRS406-gvps74(Δ197-208)-mNeon-vps74(Δ197-208)* | This study |
| SARY9048 | *MATa his3Δ1 leu2Δ0 met15Δ0 ura3Δ0 vps74::kanMX4 mnn5::MNN5-9myc-loxP-HIS3-loxP pRS406-gvps74(Δ197-208)-mNeon-PH-vps74(Δ197-208)* | This study |
| **Figure 5** |  |  |
| SARY8751 | *MATa his3Δ1 leu2Δ0 met15Δ0 ura3Δ0 vps74::kanMX4 mnn5::MNN5-9myc-loxP-HIS3-loxP* | This study |
| SARY8982 | *MATa his3Δ1 leu2Δ0 met15Δ0 ura3Δ0 vps74::kanMX4 mnn2::MNN2-9myc-loxP mnn4::MNN4-9myc-loxP transformed with gGRX7-4HA-pRS416* | This study |
| SARY9104 | *MATa his3Δ1 leu2Δ0 met15Δ0 ura3Δ0 vps74::kanMX4 dcr2::DCR2-9myc-loxP-HIS3-loxP* | This study |
| SARY9121 | *MATa his3Δ1 leu2Δ0 met15Δ0 ura3Δ0 vps74::kanMX4 grx7::GRX7-9myc-loxP-HIS3-loxP* | This study |

| **Figure 6** |  |  |
| --- | --- | --- |
| SARY8082 | *MATa his3Δ1 leu2Δ0 met15Δ0 ura3Δ0 ted1::kanMX4* transformed with *gTED1-pRS416* | This study |
| SARY8751 | *MATa his3Δ1 leu2Δ0 met15Δ0 ura3Δ0 vps74::kanMX4 mnn5::MNN5-9myc-loxP-HIS3-loxP* | This study |
| SARY8771 | *MATa his3Δ1 leu2Δ0 met15Δ0 ura3Δ0 vps74::KANMX4 mnn5::MNN5-9myc-loxP-HIS3-loxP pRS406-gVPS74* | This study |
| SARY8792 | *MATa his3Δ1 leu2Δ0 met15Δ0 ura3Δ0 vps74::mNeon-VPS74* | This study |
| Y04208 | *MATa his3Δ1 leu2Δ0 met15Δ0 ura3Δ0 vps74::KANMX4* | EUROSCARF |
| **Figure EV1** |  |  |
| SARY8627 | *MATa his3Δ1 leu2Δ0 met15Δ0 ura3Δ0 vps74::gVPS74-mNeon* | This study |
| SARY9097 | *MATa his3Δ1 leu2Δ0 met15Δ0 ura3Δ0 vps74::gVPS74-mNeon grx7::GRX7-9myc-loxP-HIS3-loxP* | This study |
| SARY9109 | *MATa his3Δ1 leu2Δ0 met15Δ0 ura3Δ0 vps74::gVPS74-mNeon mnn5::MNN5-9myc-loxP-HIS3-loxP* | This study |
| SARY9110 | *MATa his3Δ1 leu2Δ0 met15Δ0 ura3Δ0 vps74::gVPS74-mNeon och1::OCH1-9myc-loxP-HIS3-loxP* | This study |
| **Figure EV2** |  |  |
| SARY8627 | *MATa his3Δ1 leu2Δ0 met15Δ0 ura3Δ0 vps74::gVPS74-mNeon* | This study |
| SARY9011 | *MATa his3Δ1 leu2Δ0 met15Δ0 ura3Δ0 vps74::gvps74(W88A)-mNeon* | This study |
| SARY9019 | *MATa his3Δ1 leu2Δ0 met15Δ0 ura3Δ0 vps74::gvps74(R97A)-mNeon* | This study |
| SARY8598 | *MATa his3Δ1 leu2Δ0 met15Δ0 ura3Δ0 vps74::KANMX4 PRS406-gvps74(K178A, R181A)-mNeon* | This study |
| SARY9024 | *MATa his3Δ1 leu2Δ0 met15Δ0 ura3Δ0 vps74::gvps74(Δ197-208)-mNeon* | This study |
| **Figure EV3** |  |  |
| SARY8388 | *MATa his3Δ1 leu2Δ0 met15Δ0 ura3Δ0 ted1::kanMX4 vps74::loxP* transformed with *gTED1-pRS416* | This study |
| SARY8417 | *MATa his3Δ1 leu2Δ0 met15Δ0 ura3Δ0 ted1::kanMX4 vps74::loxP* transformed with *gVPS74-pRS415* | This study |
| SARY8907 | *MATa his3Δ1 leu2Δ0 met15Δ0 ura3Δ0 ted1::kanMX4 vps74::loxP* transformed with *gvps74(<sup>87</sup>AAAA<sup>90</sup>)-pRS415* | This study |
| SARY8761 | *MATa his3Δ1 leu2Δ0 met15Δ0 ura3Δ0 ted1::kanMX4 vps74::loxP* transformed with *gvps74(W88A)-pRS415* | This study |
| **Figure EV4** |  |  |
| BY4741 | *MATa his3Δ1 leu2Δ0 met15Δ0 ura3Δ0* | Lab collection |
| Y04208 | *MATa his3Δ1 leu2Δ0 met15Δ0 ura3Δ0 vps74::KANMX4* | EUROSCARF |

**Table 2.**
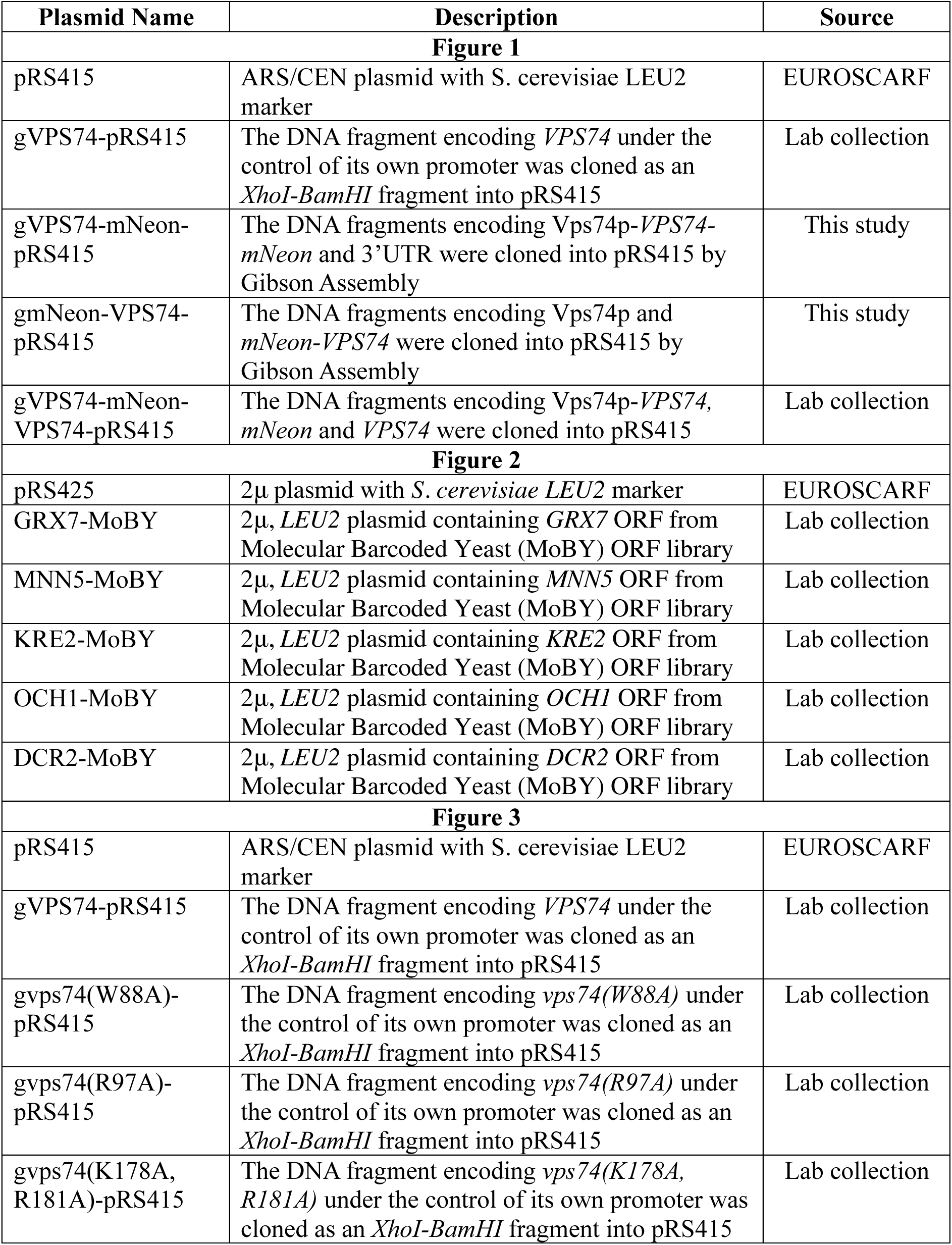
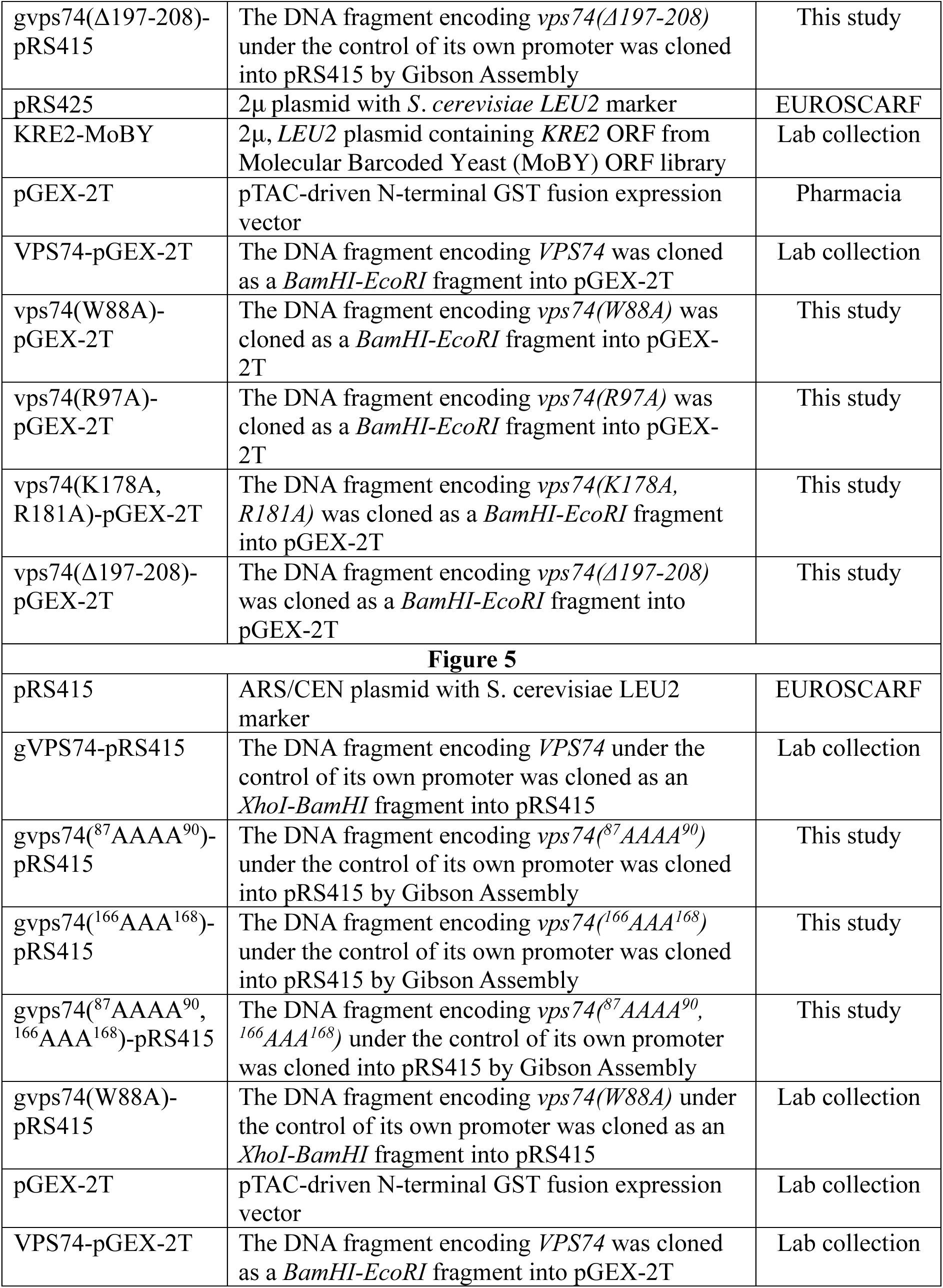
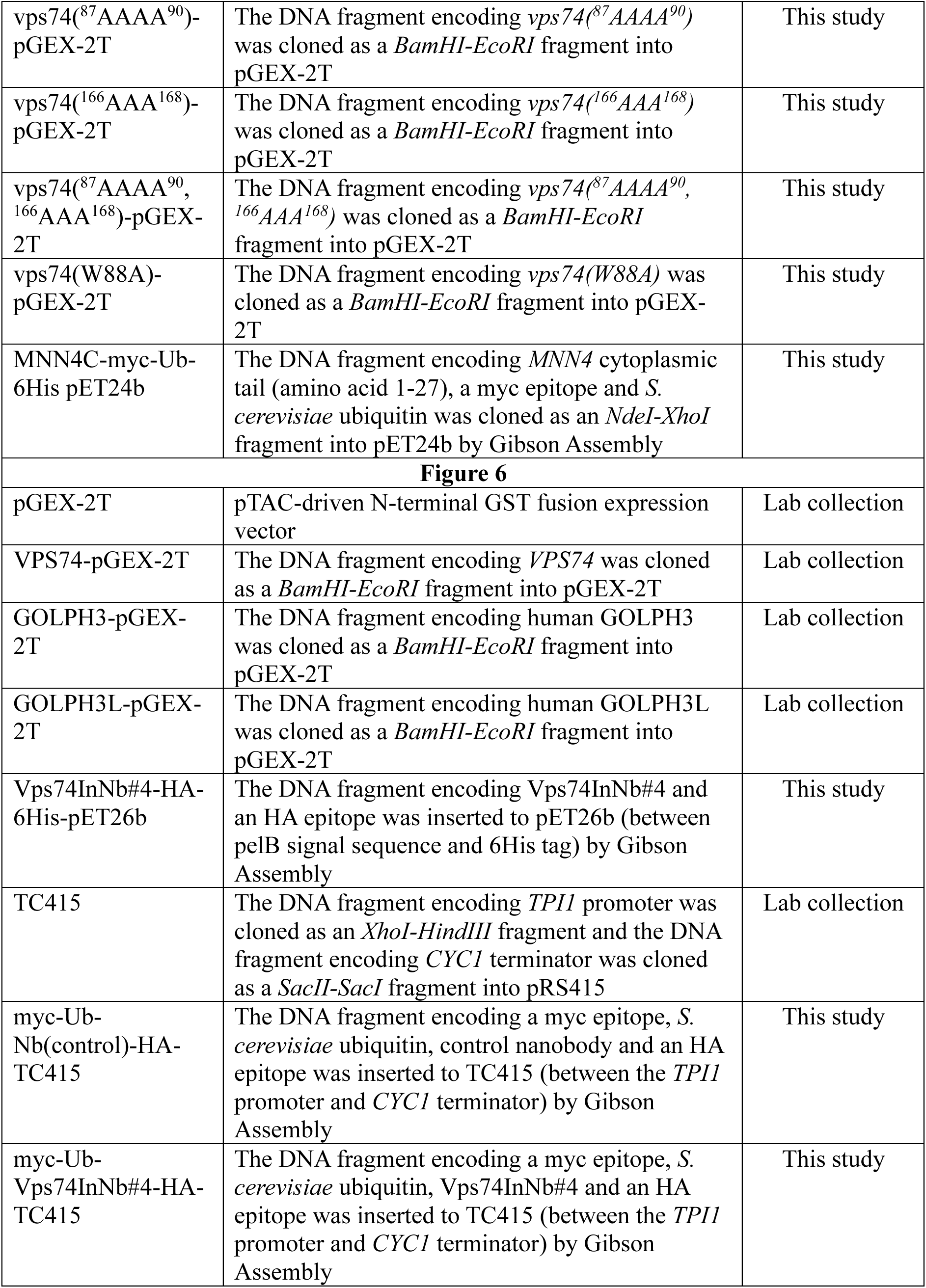
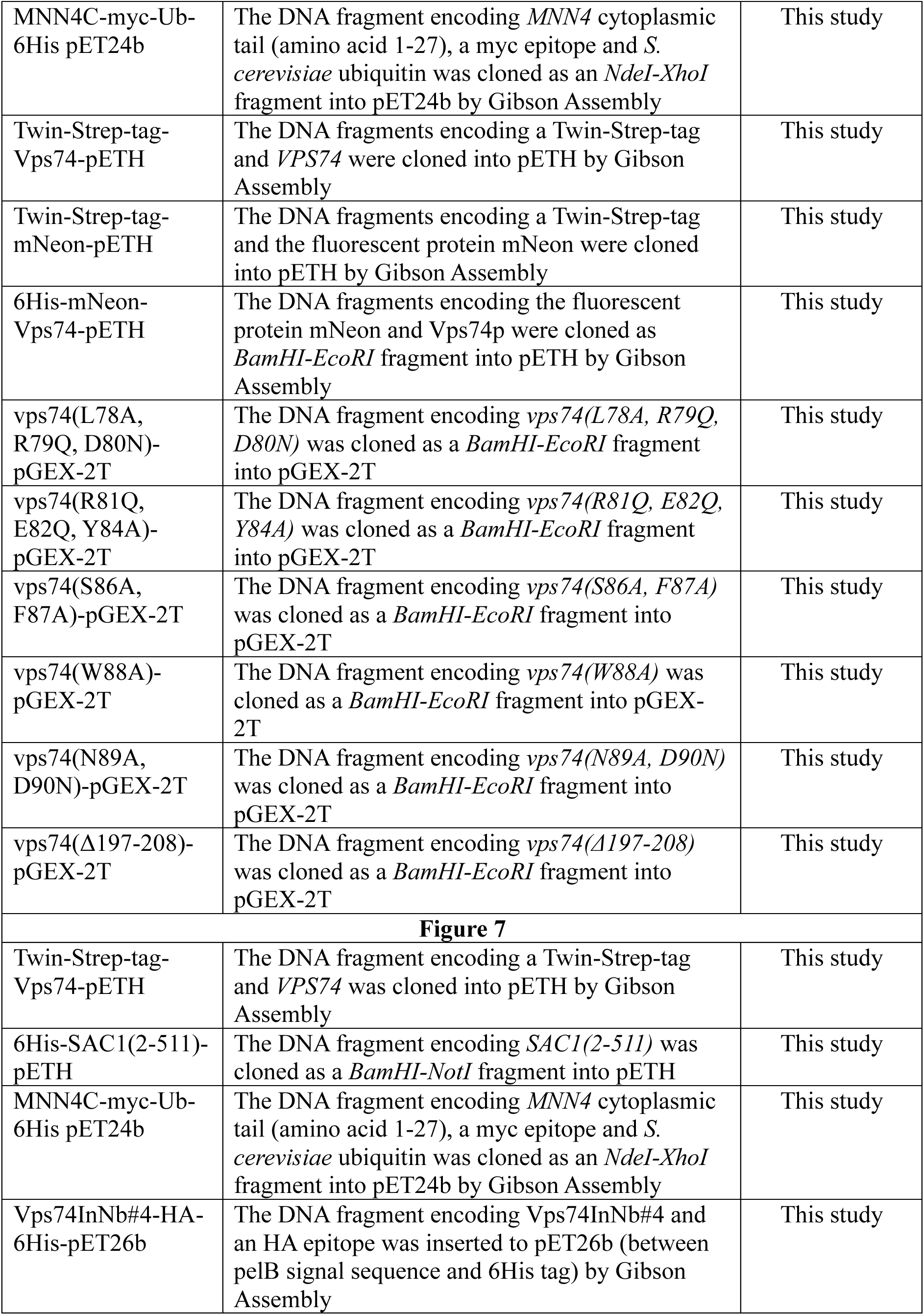
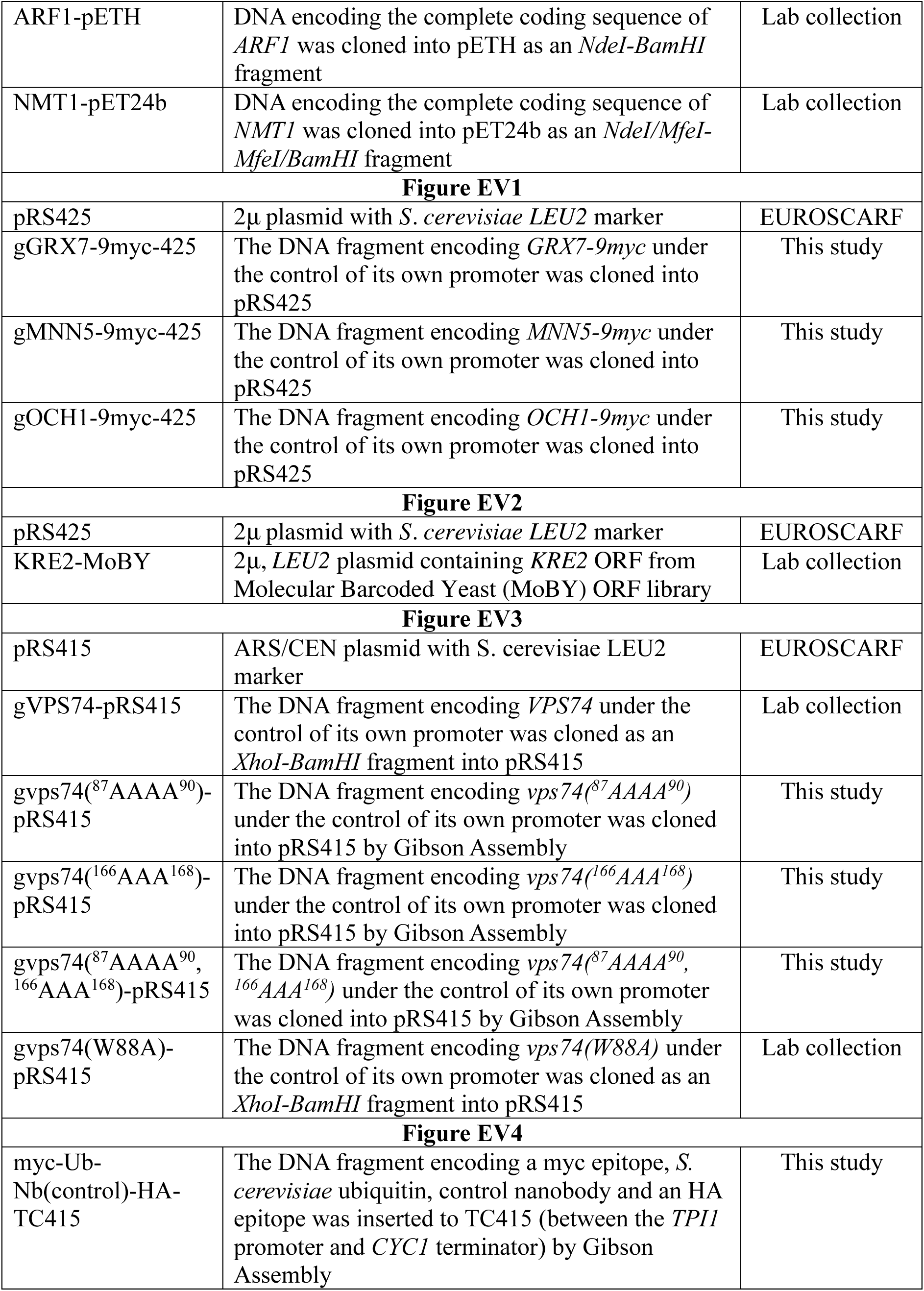
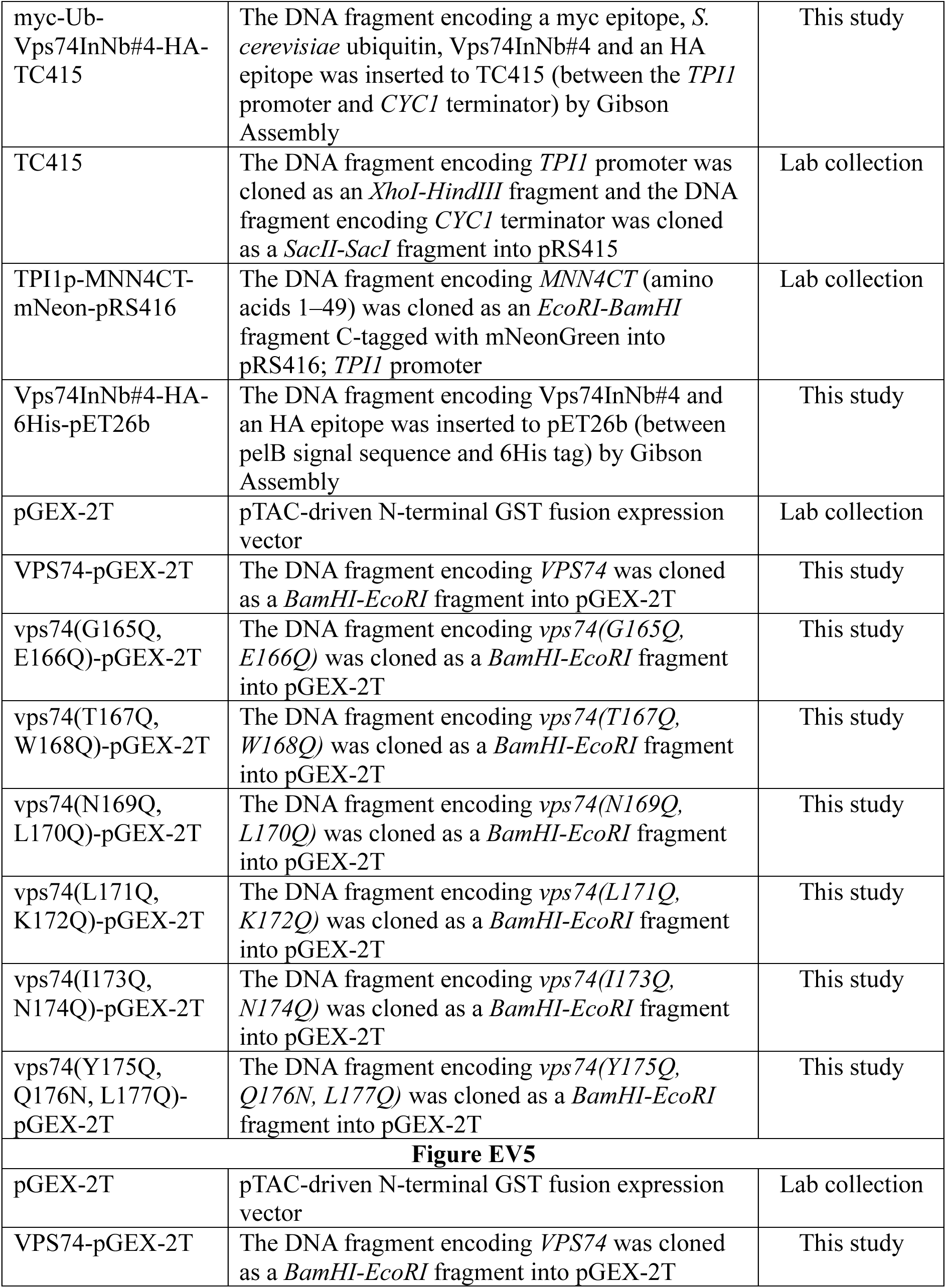
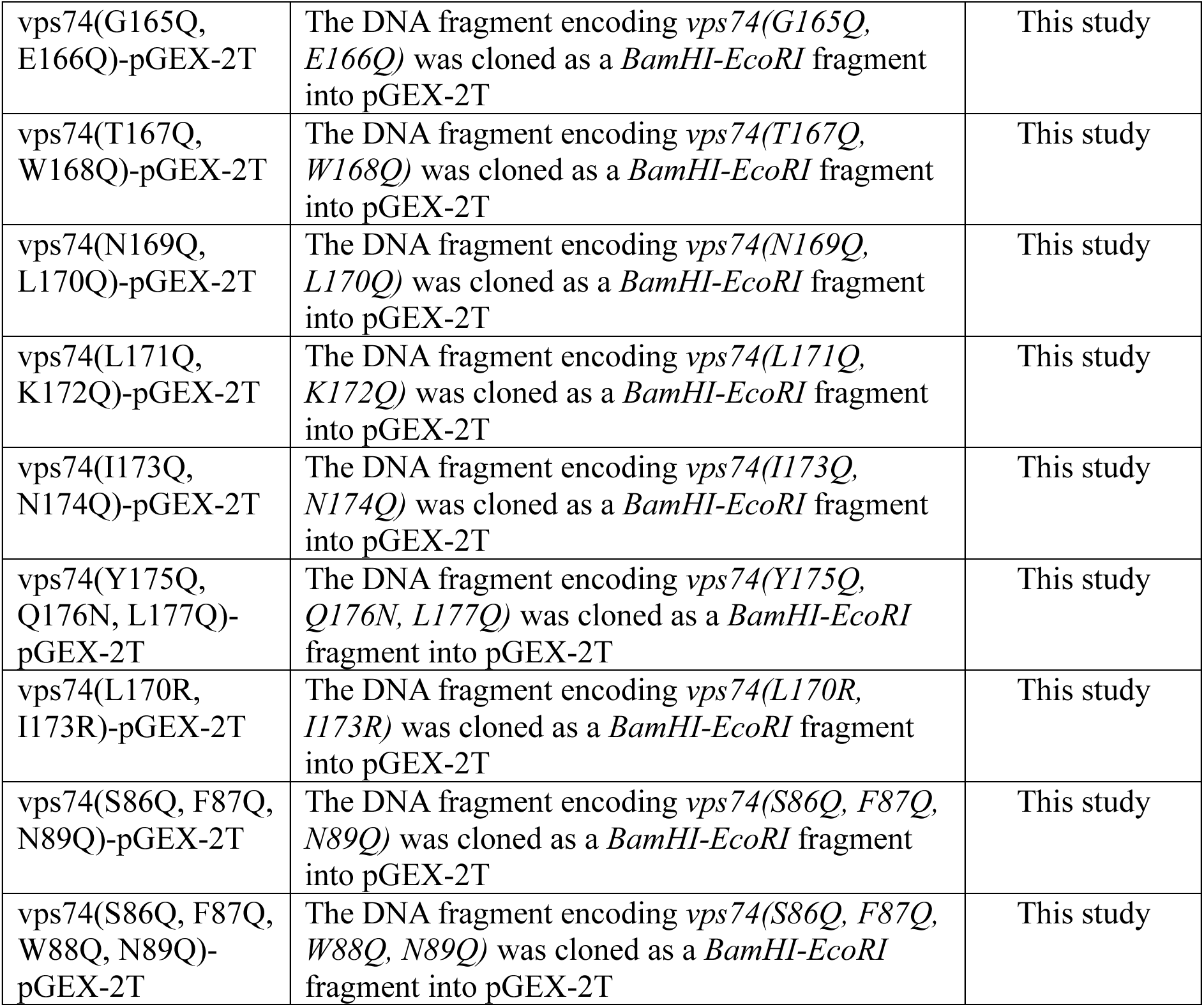
Plasmids used in this study.

**Table 3.**
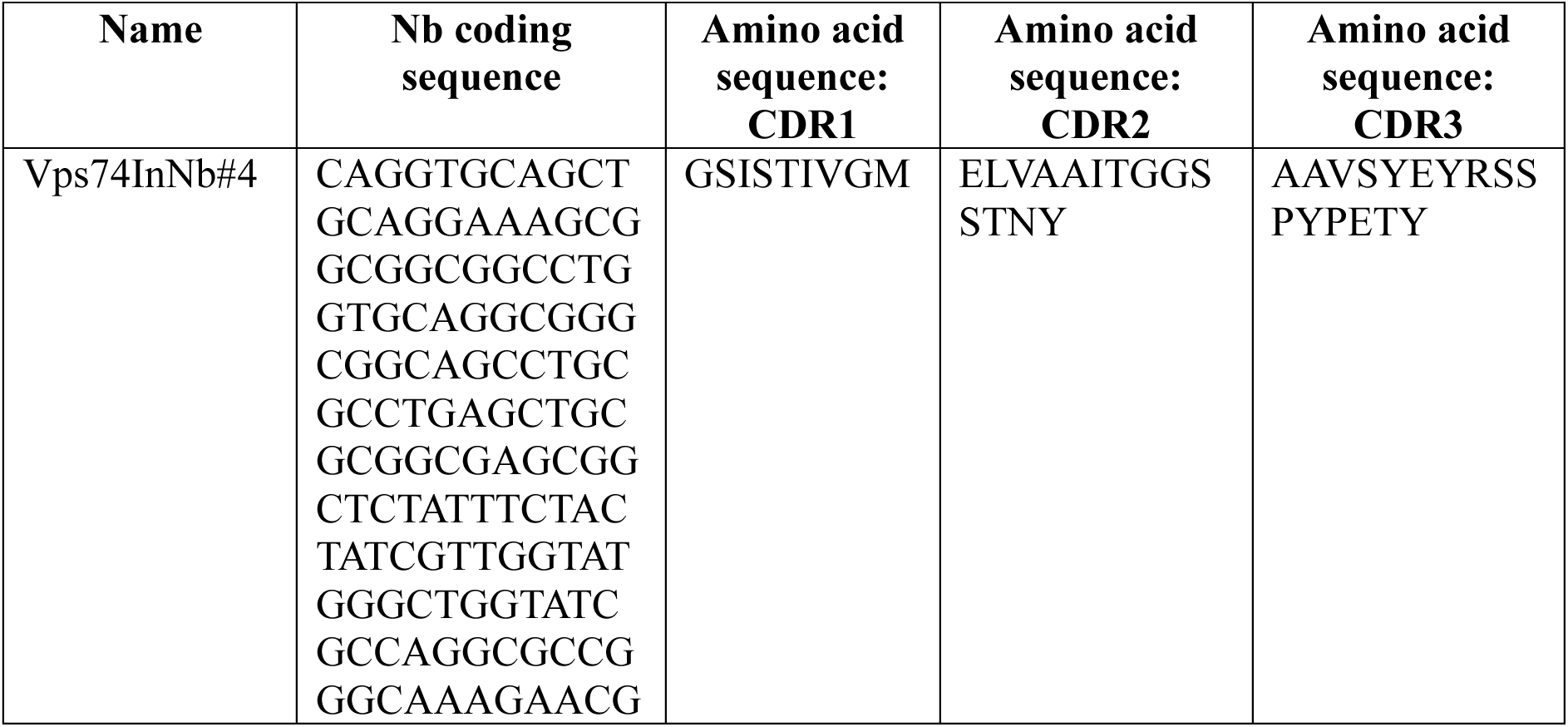
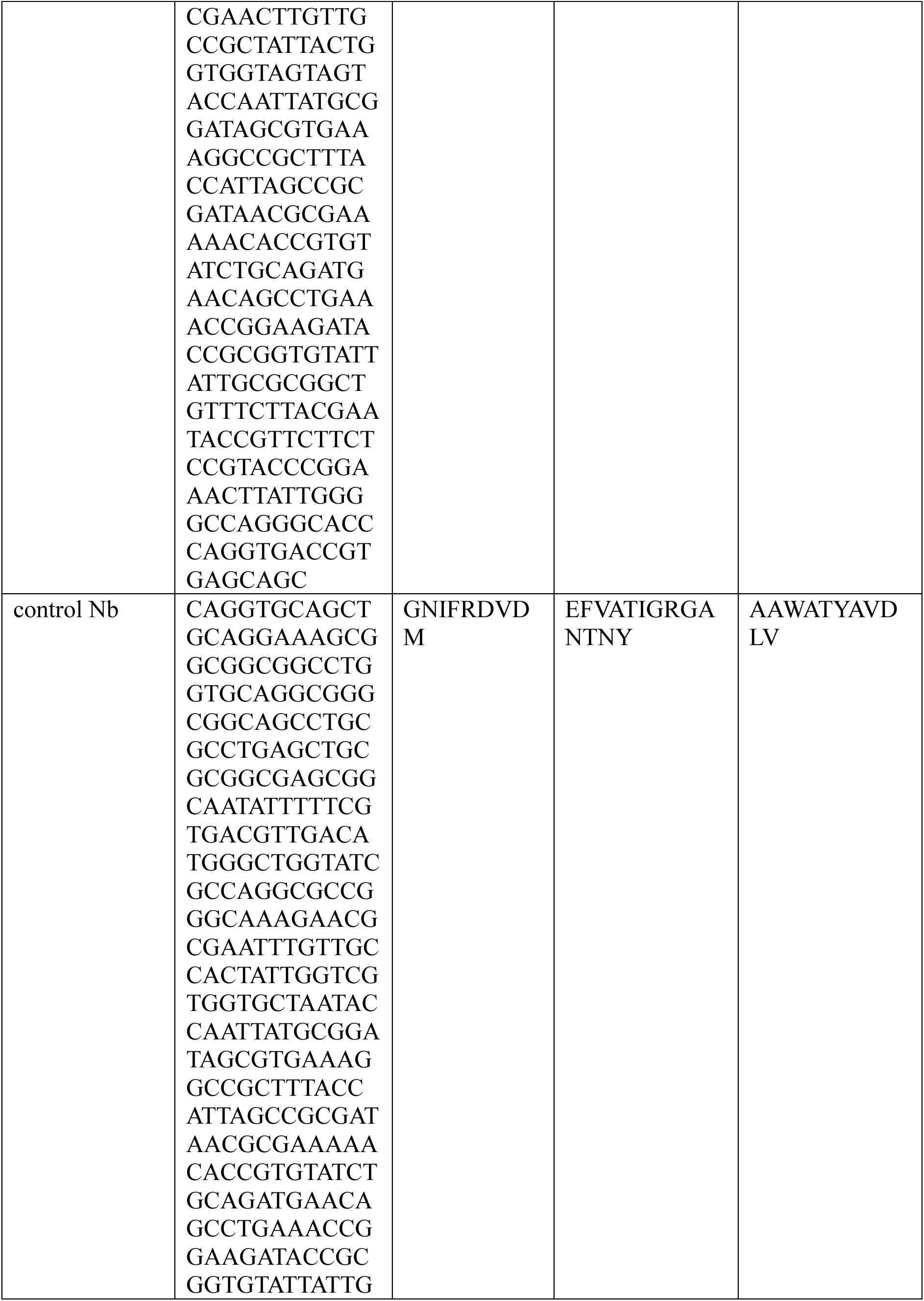
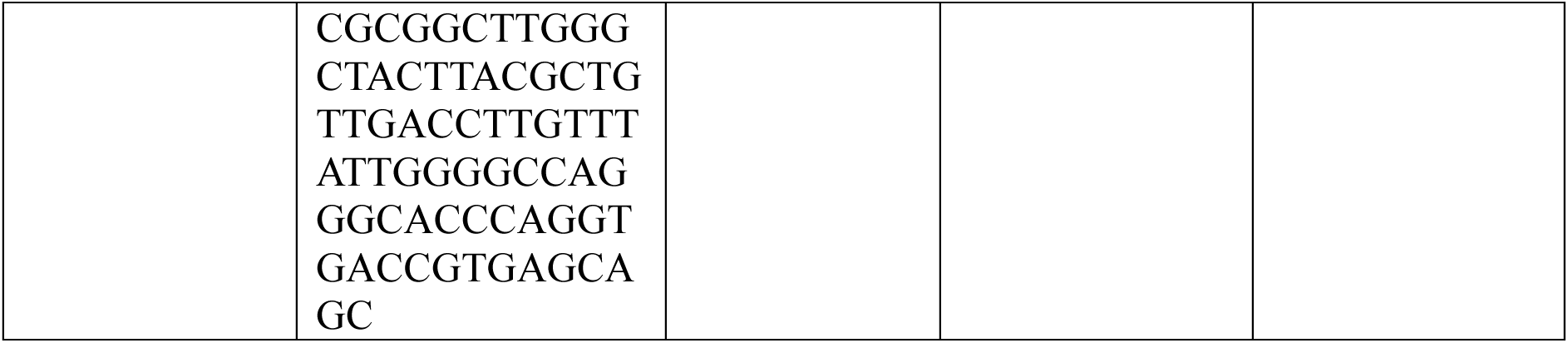
Amino acid sequence of nanobodies (Nbs) identified in this study.

**Table 4.**
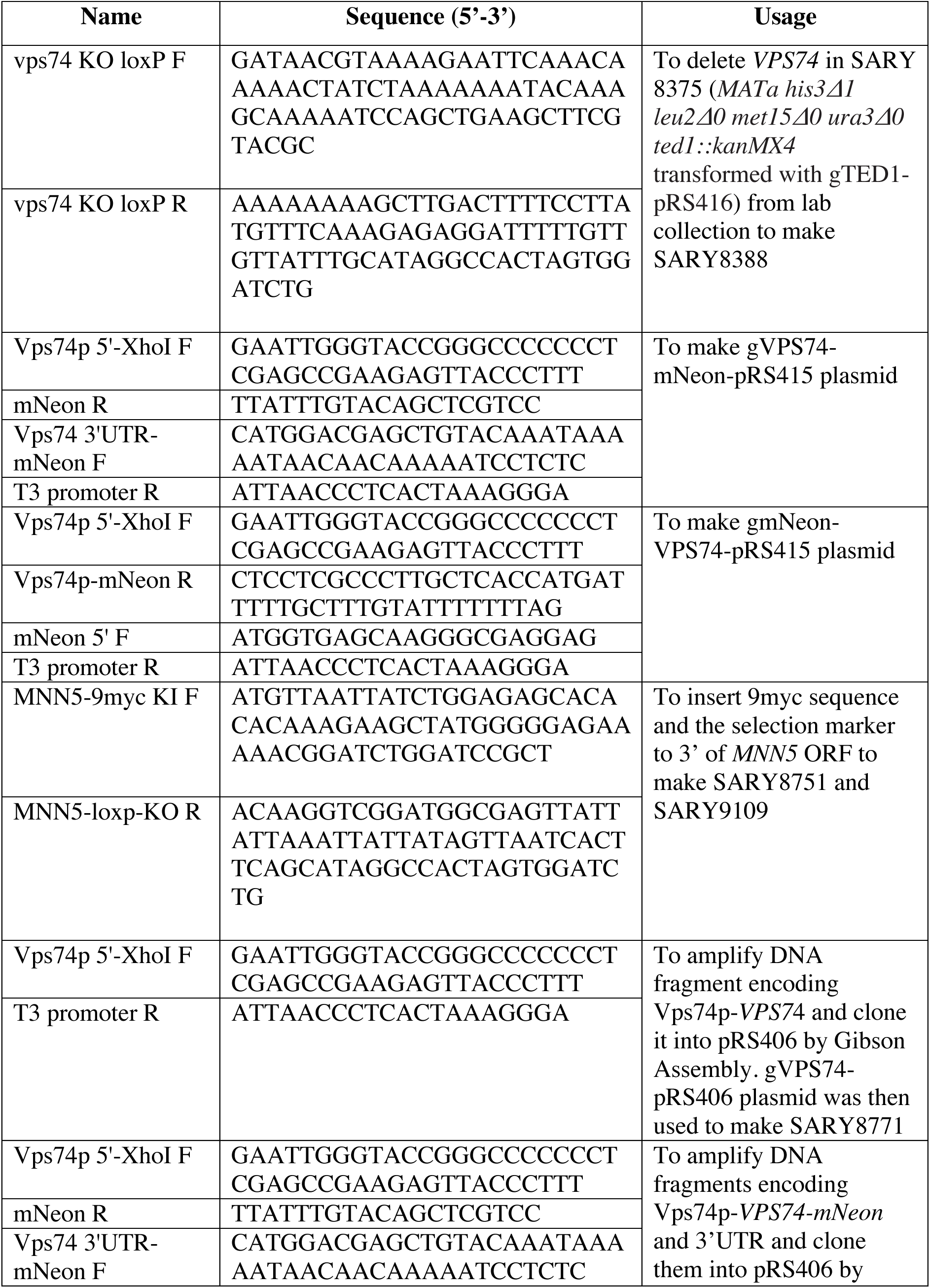
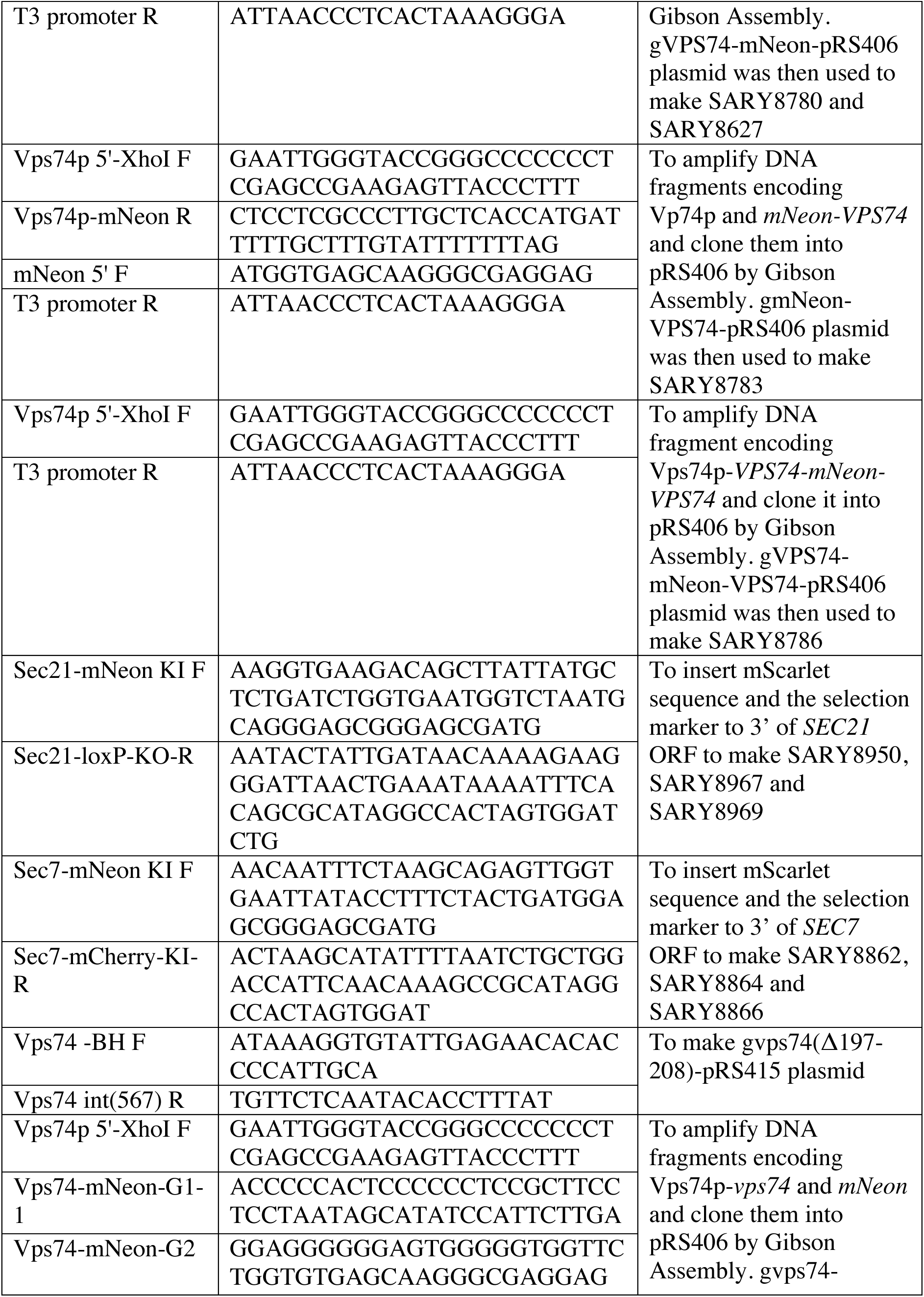
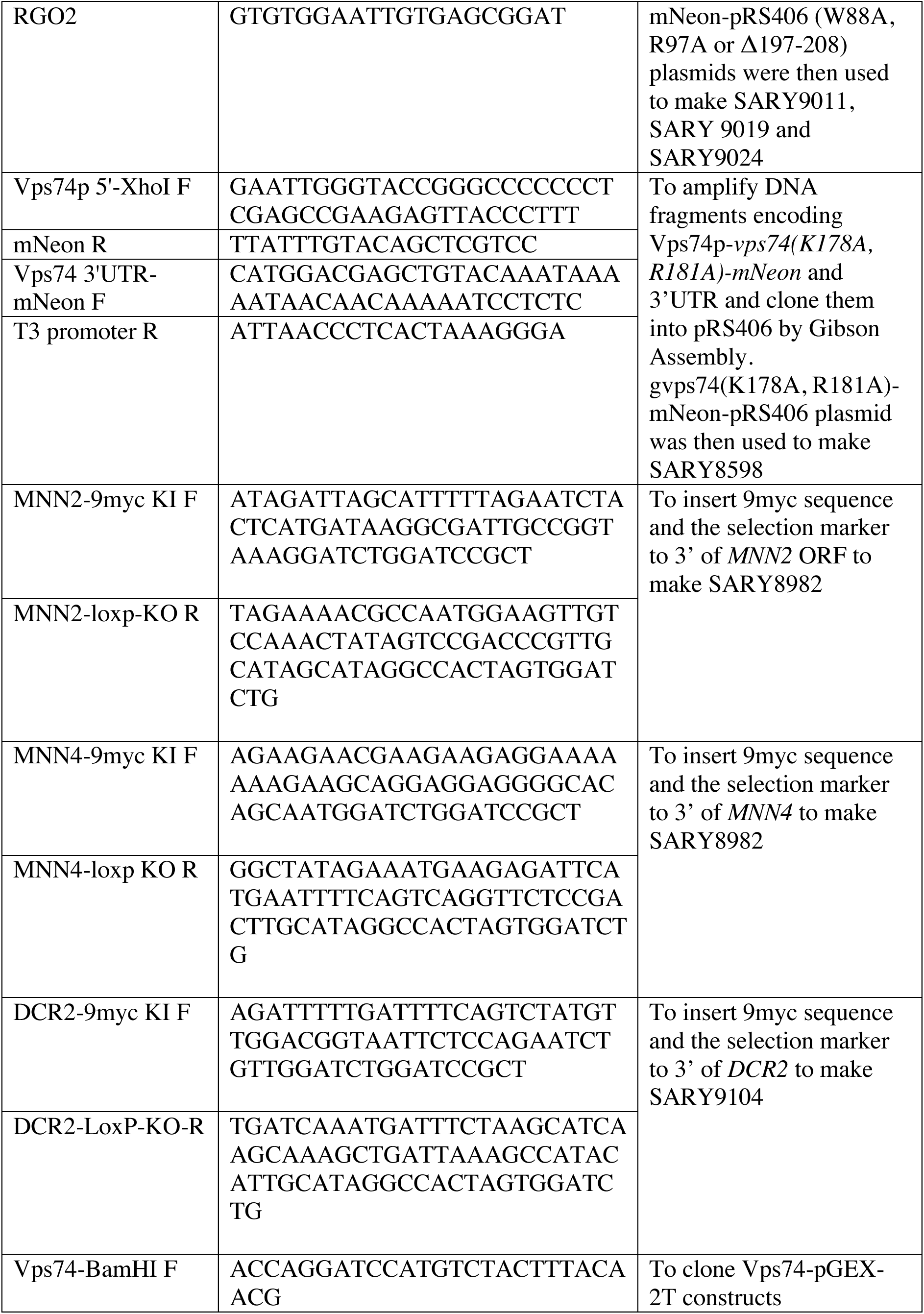
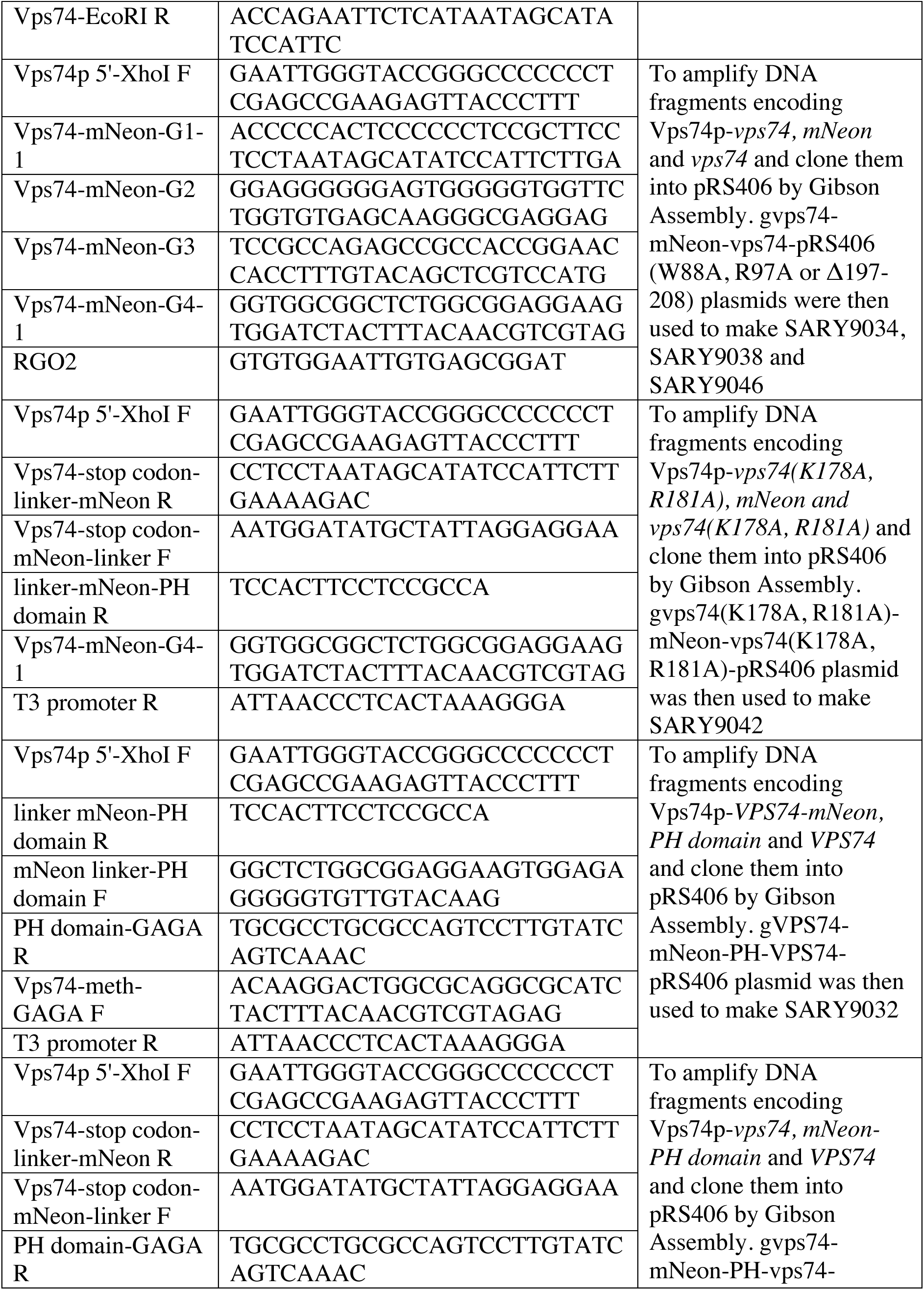
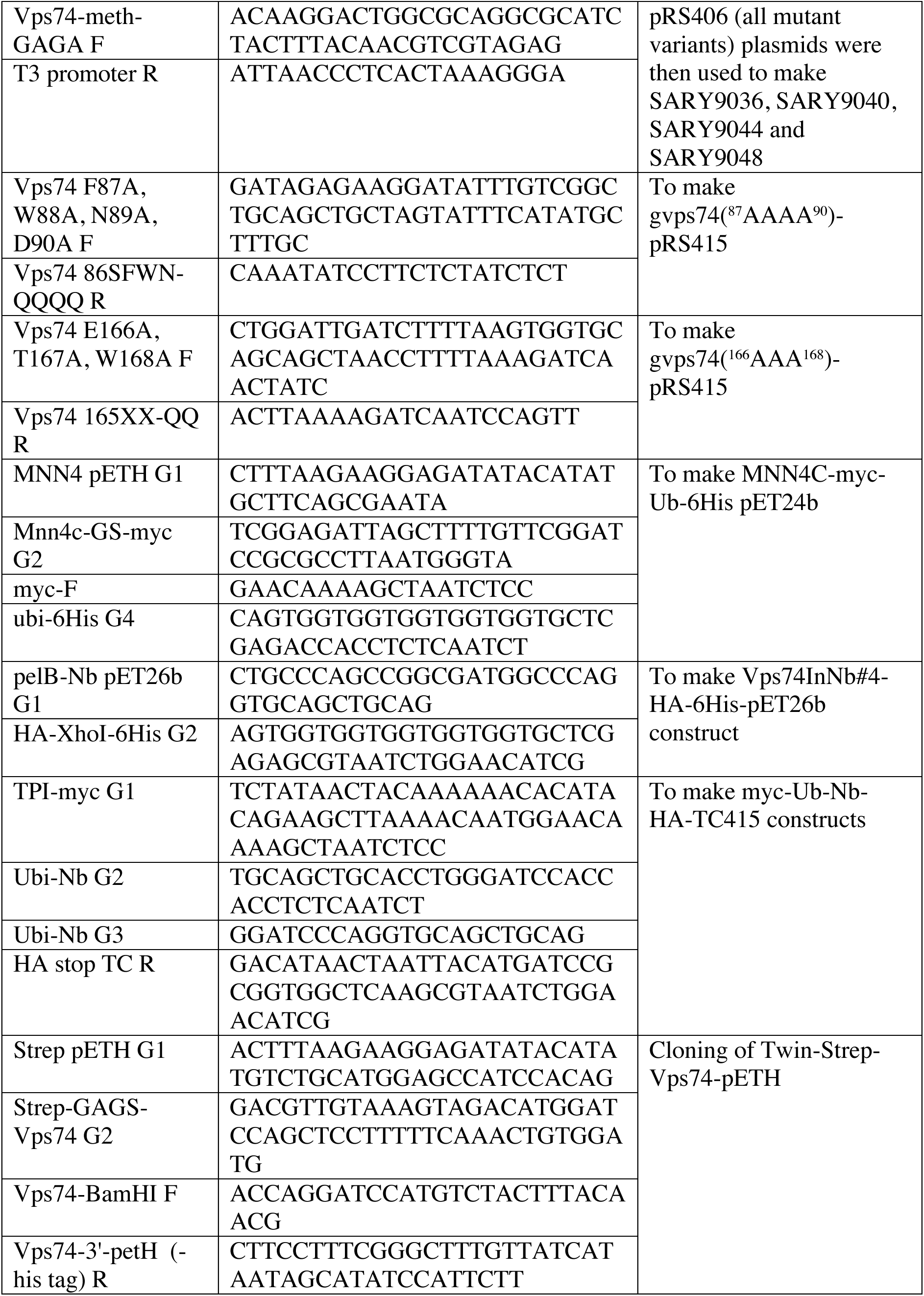
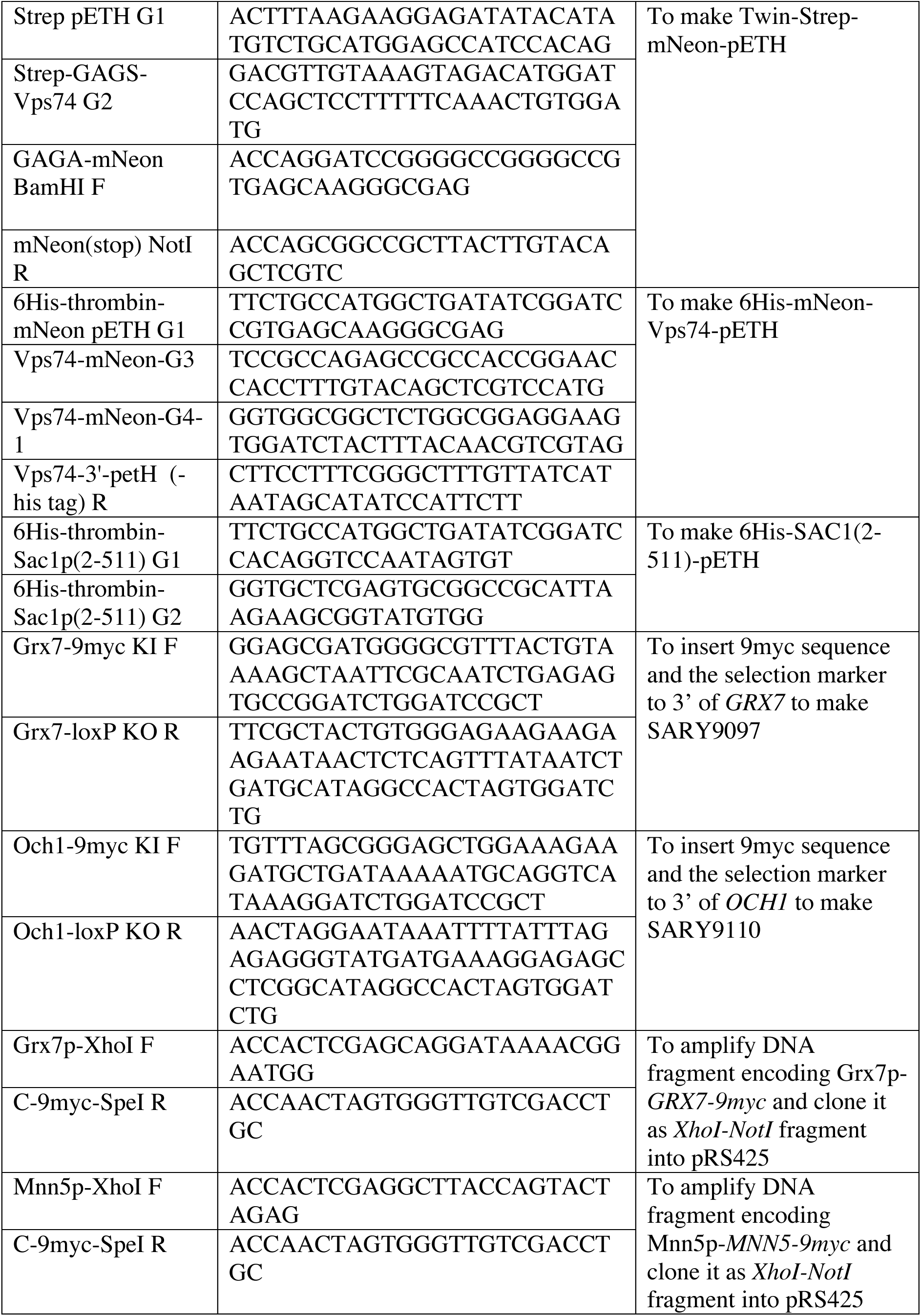
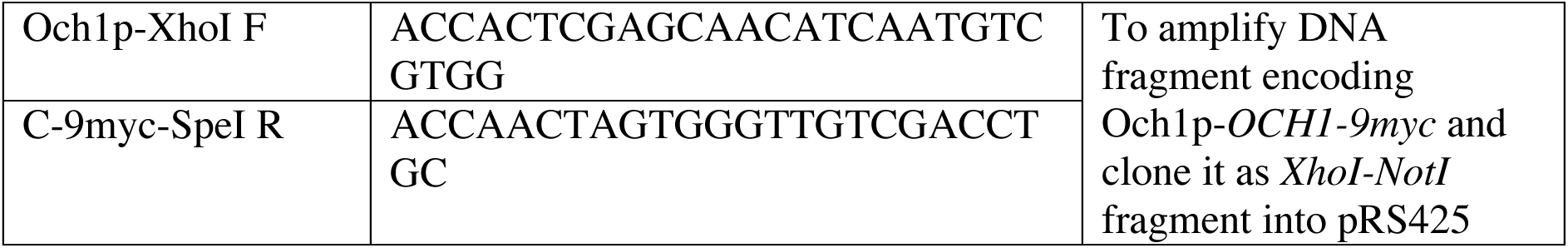
Primers used in this study.

**Table 5.** Reagents, software and equipment used in this study.

| **REAGENT or RESOURCE** | **Source** | **Identifier** |
| --- | --- | --- |
| **Antibodies** |  |  |
| Mouse monoclonal anti-myc (dilution factor 1:1000) | Roche | Cat#11667149001 |
| Mouse monoclonal anti-HA (dilution factor 1:1000) | Roche | Cat#11584000000 |
| Mouse monoclonal anti-Pgk1 (dilution factor 1:10000) | Molecular Probes | Cat#459250 |
| Rabbit polyclonal anti-Kre2 (dilution factor 1:500) | DKB lab collection | N/A |
| Rabbit polyclonal anti-GFP (dilution factor 1:4000) | Sigma | Cat#G1544 |
| Rabbit polyclonal anti-Vps74 (dilution factor 1:4000) | DKB lab collection | N/A |
| Rabbit polyclonal anti-Arf1 (dilution factor 1:2000) | DKB lab collection | N/A |
| Goat polyclonal anti-mouse IgG (dilution factor 1:3000) | Sigma | Cat#A8924 |
| Donkey polyclonal anti-rabbit IgG (dilution factor 1:3000) | GE healthcare | Cat#NA934 |
| **Chemicals, peptides, and recombinant proteins** |  |  |
| Fluorescent Brightener 28 (calcofluor white-CFW) | Sigma | Cat#F3543 |
| 5-Fluoroorotic acid (5-FOA) | Zymo Research | Cat#F9001-1 |
| Trichloroacetic acid (TCA) | Sigma | Cat#T8657 |
| Dithiothreitol (DTT) | USB | Cat#15397 |
| EDTA-free protease inhibitor cocktail | Roche | Cat#11873580001 |
| Pefabloc SC | Roche | Cat#11429876001 |
| 0.5 mm dia. Glass beads | BioSpec | Cat#11079105 |
| Endoglycosidase H (Endo H) | NEB | Cat# P0702S |
| Ponceau S | Sigma | Cat#P3504 |
| Coomassie Brilliant Blue R-250 | Bio-rad | Cat#161-0400 |
| Phosphate buffered saline (PBS) | Sigma | Cat#P3813 |
| Tween 20 | Affymetrix | Cat#T1003 |
| Amersham ECL Western Blotting Detection Reagent | Cytiva | Cat#RPN2106 |
| SuperSignal™ West Femto Maximum Sensitivity Substrate | Thermo Scientific | Cat#34096 |
| Isopropylthio- β-galactoside (IPTG) | Invitrogen | Cat#15529-019 |
| Glutathione Sepharose 4B | Cytiva | Cat#17-0756-05 |
| Ni-NTA His•Bind Resin | Millipore | Cat#70666-4 |
| Strep-tactin Sepharose | IBA Lifesciences | Cat#2-1201-010 |
| Strep-tactin Magnetic Microbeads | IBA Lifesciences | Cat#6-5510-050 |
| Triton X-100 | Sigma | Cat#X100 |
| Lysozyme | Sigma | Cat#20K0956 |
| Desthiobiotin | Sigma | Cat#D1411 |
| Imidazole | Sigma | Cat#I202 |
| Bovine serum albumin (BSA) | Sigma | Cat#A7906 |
| Galactose | Sigma | Cat#G0750 |
| Biotin | Sigma | Cat#B4501 |
| Sucrose | USB | Cat#21938 |
| PD-10 desalting columns packed with Sephadex G-25 resin | Cytiva | Cat#17085101 |
| nProteinA Sepharose 4 Fast Flow | Cytiva | Cat#17528001 |
| Guanosine 5'-[ $\beta$, γ-imido]triphosphate trisodium salt hydrate (GMP-PNP) | Sigma | Cat#G0635 |
| Guanosine 5'-diphosphate sodium salt (GDP) | Sigma | Cat#G7127 |
| Concanavalin A (ConA) | Sigma | Cat#C7275 |
| FM4-64 Dye | Invitrogen | Cat#T13320 |
| Concanavalin A (C7275) (ConA) | Sigma | Cat#C7275 |
| **Bacterial and virus strains** |  |  |
| *E. coli* BL21(DE3) | Lab collection | N/A |
| **Experimental models: Organisms/strains** |  |  |
| *S. cerevisiae* | Lab collection and this study | Table 1 |
| **Software and algorithms** |  |  |
| ImageJ | NIH | https://imagej.net/ij/ |
| Prism 10 | GraphPad | https://www.graphpad.com/features |
| Fiji | NIH | https://fiji.sc |
| ChimeraX | UCSF | https://www.rbvi.ucsf.edu/chimerax |
| **Other** |  |  |
| Yeast-Display Nanobody Library (NbLib) | From the laboratory of Andrew C. Kruse, PhD, Harvard University. Distributed by Kerafast. | Cat#EF0014-FP |
| StrepMan Magnet | IBA Lifesciences | Cat#6-5650-065 |
| Econo-Column Chromatography Columns, 1.0 × 10 cm | Bio-rad | Cat#7371012 |
| Q125 sonicator | QSONICA | Cat#Q125-110 |
| ChemiDoc Imaging System | Bio-rad | Cat#12003153 |

5 × 10^6^ mid-log phase *ted1Δ* cells balanced with the *TED1-URA3* plasmid (SARY8082, see Table 1) were co-transformed with 50 ng the TPI1p-CYC1t pRS415 (see Table 2) plasmid digested with *HindIII* and *NotI*, and 50 ng of purified myc-ubiquitin-nanobody-HA fragment. Transformants were selected on SD plates (-Leu) and single colonies were patched on SD plates (-Leu) with and without 5-FOA. Cells that failed to grow on SD containing 5-FOA plates were recovered from SD (-Leu) plates. Nanobody coding sequences were amplified by the PCR using the primers pelB-Nb pET26b G1 and HA-XhoI-6His G2 (see Table 4) and thereafter the fragment was cloned into the *E. coli* periplasmic expression vector pET26b, by Gibson Assembly (Gibson et al., 2009)

#### Expression and purification of nanobodies from the periplasm of bacteria

*E. coli* BL21(DE3) cells were transformed with pET26b plasmids (see Table 2) containing nanobody coding sequences. A single colony was used to inoculate the start culture in Terrific Broth (TB) and the culture was shaken at 37°C and 210 rpm overnight. The overnight culture was diluted 1:40 with fresh TB and grown under the same conditions. When the OD_600_ reached 0.6-0.8, the culture was shifted to 25°C, and 1 mM IPTG was added to induce nanobody protein synthesis. After overnight incubation at 210 rpm, cells from 100 ml culture were collected by centrifugation at 4,200 rcf for 10 min and gently resuspended in 3 ml TSE buffer (0.2 M Tris, 0.5 M sucrose, 0.5 mM EDTA, pH 8.0). The resuspended cells were incubated at room temperature for 10 min with occasional shaking. 3 ml ice-cold ddH_2_O was added to trigger osmotic shock, and the mixture was incubated on ice for 10 min with occasional shaking. The lysate was supplemented with 150 mM NaCl, 2 mM MgCl_2_ and 20 mM imidazole, and the released periplasmic extract containing nanobodies was cleared by centrifugation at 16,000 rcf for 10 min at 4°C. The periplasmic extract was added to 0.5 ml Ni-NTA beads and mixed for 1 h at 4°C. The beads were washed with 3 ml wash buffer (20 mM HEPES, 500 mM NaCl, 20 mM imidazole, pH 7.5) three times. Nanobodies were eluted with 500 μl PBS + 250 mM imidazole five times and the elution buffer was exchanged for PBS + 10% glycerol using PD-10 desalting columns containing Sephadex G-25 resin.

#### Co-immunoprecipitation assay

50 × 10^7^ mid-log phase wild-type yeast cells expressing Nb-HA were collected by centrifugation. Cells were resuspended in 1 ml lysis buffer (PBS supplemented with 1x protease inhibitor cocktail, 1 mM Pefabloc SC, 0.5 mM EDTA and 0.1% Triton X-100) and lysed with 500 μl glass beads in a bead-beater for 5 min at 4°C (1 min on, 1 min off). Yeast lysates were cleared by centrifugation at 16,000 rcf for 10 min at 4°C. A 20 μl slurry of Protein A Sepharose beads, equilibrated with lysis buffer, was mixed with yeast cell lysates for 1 h, at 4°C. Following incubation, protein-bound beads were washed with 200 μl lysis buffer three times and boiled in 50 μl SDS sample buffer supplemented with 5 mM DTT for 5 min. Bound proteins eluted from beads were analyzed by SDS-PAGE and immunoblotting.

#### *In vitro* mixing and competition assays

Bacterial lysates containing 10 ug GST-fusion proteins were mixed with 20 μl Glutathione Sepharose 4B beads equilibrated with PBS + 0.1% Triton X-100 for 1 h, at 4°C. Unbound proteins were washed away with 200 μl PBS + 0.1% Triton X-100 three times, and equimolar amounts of purified proteins of interest diluted in 100 μl PBS + 0.1% Triton X-100 were mixed with beads for 1.5 h at 4°C. Beads were washed with 200 μl PBS + 0.1% Triton X-100 three times and boiled in 30 μl SDS sample buffer supplemented with 5 mM DTT for 5 min. Bound proteins eluted from beads were analyzed by SDS-PAGE and immunoblotting.

For protein binding competition assays, 15 μl Strep-tactin Sepharose beads were bound to 10 ug of Twin-Strep-Vps74 as described above. A two-molar excess of the first protein of interest (Sac1 or Vps74InNb#4) diluted in 100 μl PBS + 0.1% Triton X-100 was mixed with loaded beads for 1h at 4°C and beads were washed with 200 μl PBS + 0.1% Triton X-100 three times. Equimolar amounts of the second protein of interest diluted in 100 μl PBS + 0.1% Triton X-100 was then mixed with Twin-Strep-Vps74 bound beads for 1.5 h at 4°C. After incubation, beads were washed with 200 μl PBS + 0.1% Triton X-100 three times and boiled in 30 μl SDS sample buffer supplemented with 5 mM DTT for 5 min. Bound proteins eluted from beads were analyzed by SDS-PAGE and immunoblotting.

For *in vitro* assays using purified N-myristoylated Arf1, 2 μg Arf1 was incubated in 20 μl Arf1 loading buffer (20 mM HEPES, 100 mM NaCl, 1 mM EDTA, 1 mM MgCl_2_, 5 mg/L BSA, 0.1% Triton X-100, pH 7.2) supplemented with 40 μM GMP-PNP for 1.5 h at 32°C. Arf1 pre-loaded with nucleotides was then used in the *in vitro* assays.

#### Quantification of Coomassie Blue staining, Ponceau S staining, and immunoblots

ImageJ (Schneider et al., 2012) was used to measure the integrated intensities of protein bands from images captured using a ChemiDoc Imaging System. For measurements of steady-state levels of proteins from yeast whole cell extracts, the intensities of proteins of interest were normalized to the corresponding intensities of Pgk1. Intensities in the wild-type group were defined as 100%, and the intensities of other samples were normalized and expressed as percentages thereof accordingly. For the *in vitro* mixing and binding competition assays, the intensities of proteins of interest were normalized to the corresponding intensities of Vps74. Then the average intensity of all samples in the wild-type group or “no competition” group was defined as 100%.

#### Staining of the yeast vacuole limiting membrane

Yeast cells were grown to the early-mid log phase (OD_660_ ∼0.5 - 0.8) in SD media and cells from 0.5 ml of culture media were harvested and resuspended in 50 μl SD media containing 20 μM FM 4-64 dye. After incubation at 30°C for 1 h, cells were washed with 200 μl media to remove excess dye and resuspended in 1 ml media. Cells were grown at 25°C for 2 h, and thereafter examined by fluorescent microscopy.

#### Fluorescence microscopy

Yeast knock-in strains or transformants were grown to the early-mid log phase (OD_660_ ∼0.5-0.8) in SD media and thereafter cells were collected by centrifugation. ∼10^7^ cells were resuspended in 100 μl SD media and 0.8 μl of the cell suspension was applied onto a ConA-coated (200 μg/mL) glass slide (Hendley-Essex, UK). A coverslip was added on top of the slide and immobilized with nail polish. For Figure 6 and Figure EV4, samples were visualized with a Nikon Eclipse 80i microscope (Nikon instruments, Japan) equipped with a SPOR-RT3 monochrome w/o IR camera (Diagnostic Instruments, inc. Sterling Heights, MI, USA). Cells were photographed through a Nikon Plan Apo VC 100X/1.4 oil immersion objective lens. For all other Figures, images were obtained with a Zeiss LSM 980 Confocal Microscope (Carl Zeiss Microscopy GmbH, Germany) and a 100X/1.4 oil immersion objective lens. For co-localization studies fluorescent signals were acquired simultaneously. Images were processed by Fiji (Schindelin et al., 2012).

#### Quantification of Vps74 puncta

Vps74 fluorescence puncta were quantified using the 3D-Objects Counter plug-in in Fiji software (Bolte et al., 2006). In-focus cells were chosen from a bright field image to minimize measurement bias. Next, a threshold was manually applied to a fluorescence image to remove cytosolic signals from cells and isolate isotropic, round-shaped structures comprised of 5 or more pixels. Such identified “objects” were further delineated by performing connexity analysis, which allowed for the segmentation of individualized puncta. Corrections were made if necessary, following the visual inspection. Pixels with fluorescence intensity values that exceeded the threshold but did not resemble an oval-like shape, were excluded from the analysis. Finally, geometrical centers of the obtained foci were established automatically, and their number was counted. Multiple fields of view with 20 cells each were examined for every transformant.

#### Quantification of Vps74 puncta co-localizing with *cis*, medial and *trans* Golgi proteins

Co-localization analysis were repeated 3 times (n=3) and at least 20 cells were assessed for each technical replicate. Cells were subjected to object-based co-localization analysis using the Just Another Co-localization Plug-in (JACoP) implemented within Fiji software (Bolte et al., 2006). The degree of colocalization was determined by the percentage overlap of the geometrical centers of Vps74 puncta falling inside segmented objects of the red channel (i.e., particles of the respective Golgi markers: Sed5, Sec21, Sec7).

#### Statistical analysis

Where only two data groups were tested, data were analysed using unpaired t-tests assuming equal variance. For data with more than two groups, one-way ANOVA followed by Tukey’s or Dunnett’s multiple comparison tests were performed. For data sets with two independent variables, two-way ANOVA followed by Tukey’s or Sidah’s multiple comparison tests were performed. Individual data points, means and s.d. are presented on graphs together with P values. Statistical analysis and graph generation were performed using Prism 10 software (GraphPad, USA).

## RESULTS

### Vps74 localizes to multiple Golgi *cisternae*

Vps74 and its homologs (the GOLPH3) are widely distributed throughout eukaryotes. These proteins are peripheral Golgi membrane proteins that reportedly associate with membranes through their PI4P-binding site and / or through a conserved β hairpin (Cai et al., 2014; Wood et al., 2009). GOLPH3s function as COPI-coatomer adaptors that bind to the cytoplasmically exposed N-termini of numerous Golgi-resident integral membrane proteins (Schmitz et al., 2008; Tu et al., 2008; Welch et al., 2021). Hereafter we refer to these integral membrane proteins as clients. In addition to its client-binding ability, Vps74 reportedly stimulates the PI4P phosphatase activity of Sac1 (Cai et al., 2014; Wood et al., 2012). Presumably, where Vps74 and Sac1 distributions overlap the prevalence of PI4P is likely to be reduced (Cai et al., 2014). Given that PI4P is found predominantly in the *trans* Golgi (Audhya et al., 2000; Hama et al., 1999; Walch-Solimena and Novick, 1999), but Vps74’s clients are distributed throughout the Golgi, how does Vps74 bind to Golgi membranes with comparatively little PI4P?

To begin to address this question, we generated Vps74 fusions to mNeon to visualize the protein in cells. Although others have shown that N-terminally GFP-tagged Vps74 localizes to Golgi-like puncta, a more robust test of the functional consequences of fusing a fluorescent protein to Vps74 is needed (Schmitz et al., 2008; Sardana et al., 2021). For this purpose, we generated three mNeon-tagged forms of Vps74 (Figure 1A). All three mNeon / Vps74 fusion proteins were functional as judged by their ability to support the synthetic lethal growth phenotype of cells lacking the *VPS74* and *TED1* genes (Chen et al., 2021) (Figure 1B). Similarly, *vps7411 ted111* cells expressing the mNeon / Vps74 fusion proteins grew like control cells at 37°C (Figure 1C). Although the expression levels of Vps74, Vps74-mNeon and mNeon-Vps74 were similar (Figure 1D), none of the Vps74 / mNeon constructs were as effective as WT Vps74 in maintaining the steady-state levels of Dcr2 (Figure 1E). However, in the case of Mnn5 and Kre2, mNeon-Vps74 was more effective in the retention of these clients than Vps74-mNeon (Figure 1F) (Tu et al., 2008; Wang et al., 2020). Interestingly, the tandem chimera (Vps74-mNeon-Vps74) was no more effective than its monomeric counterparts in the retention of Dcr2, Mnn5 and Kre2 (Figure 1E and F) suggesting that tethering two molecules of Vps74 did not increase the effectiveness of the adaptor.

To establish the extent to which the three tagged forms of Vps74 could bind to different Golgi cisternae, we integrated the various mNeon-tagged variants at the *VPS74* locus in cells expressing mCherry-Sed5 (a *cis* Golgi resident), Sec21-mScarlet (a *cis*, medial and *trans* Golgi resident) or Sec7-mScarlet (a *trans* Golgi resident) (Figure 1G). Vps74-mNeon showed the fewest puncta ∼ 5 puncta per 20 cells, whereas mNeon-Vps74 and Vps74-mNeon-Vps74 showed roughly the same number of puncta per cell (∼35 - 40 per 20 cells) (Figure 1H). Despite the reduction in total puncta, the distribution of Vps74-mNeon across Golgi cisternae, as judged by colocalization with Sed5, Sec21 and Sec7, was like that seen with mNeon-Vps74 and Vps74-mNeon-Vps74 (Figure 1I). These data establish that Vps74 is distributed more or less equally across the *cis*, medial and *trans* cisternae. The relative decrease in Vps74-mNeon Golgi puncta is in accord with our findings that cells expressing this version of the adaptor show a reduction in the steady-state levels of Dcr2, Mnn5 and Kre2 (Figure 1E and F). We conclude that Vps74-mNeon is functional but hypomorphic due to the protein’s reduced capacity to bind to Golgi membranes. Importantly, the recruitment of Vps74 to multiple cisternae correlates well with the distribution of the adaptor’s clients in cells (Tojima et al., 2024; Wang et al., 2020).

The co-localization data presented in Figure1G, H, and I showed that the three Vps74 mNeon fusion proteins colocalize with ∼ 40 - 45% of Sed5 (*cis* Golgi) positive puncta, with ∼30 - 35% of Sec7 (*trans* Golgi) positive puncta and with ∼75 – 90 % of Sec21 (*cis*, medial and *trans*). These findings reveal that irrespective of the positioning of mNeon on Vps74 all three variants display similar Golgi distributions. Apparently, Vps74 can bind to membranes with lower steady-state levels of PI4P than found in the *trans* Golgi, implying that an additional / alternative mechanism may account for the protein’s localization to *cis* and medial cisternae (Cai et al., 2014; Wood et al., 2009).

### Client over-expression is sufficient to recruit Vps74 to Golgi membranes

To explore the prospect of an additional membrane-binding mechanism, we asked if Vps74 could be recruited to the Golgi directly by binding to its clients. To address this, we introduced high copy number plasmids containing various client genes into cells expressing Vps74-mNeon as their sole source of the adaptor. We chose Vps74-mNeon for these experiments as the fusion protein, whilst being functional (Figure 1), was found on fewer Golgi puncta, making any increase more readily apparent. We selected a subset of Vps74 clients to test as well as the non-Vps74 client *OCH1* (Wang et al., 2020). The data presented in Figure 2A reveal that when transformed with high copy number plasmids containing the client genes *MNN5*, *KRE2* (Tu et al., 2008; Wang et al., 2020) and *GRX7*, cells had a larger number of Vps74-mNeon puncta than those containing empty vector or the non-Vps74 client *OCH1* plasmid. On average, the presence of high copy number plasmids expressing client genes increased puncta to ∼ 20 per 20 cells (or 4-fold) compared to the non-client containing plasmid or vector only which showed ∼ 5 puncta per 20 cells (Figure 2A). The steady-state levels of Grx7, Mnn5, Kre2 and Och1 were indeed increased (∼ 5 – 10-fold) relative to wildtype cells as judged by immunoblots of whole cell extracts from cells transformed with the high copy number plasmids in which clients (excluding Kre2) were expressed as C-terminal fusion proteins to 9 copies of the myc epitope tag (Figure EV1A). The addition of an epitope tag to these clients did not affect the capacity of over-expressed clients to increase the number of Vps74-mNeon puncta (Figure EV1B)

**Figure 2.**
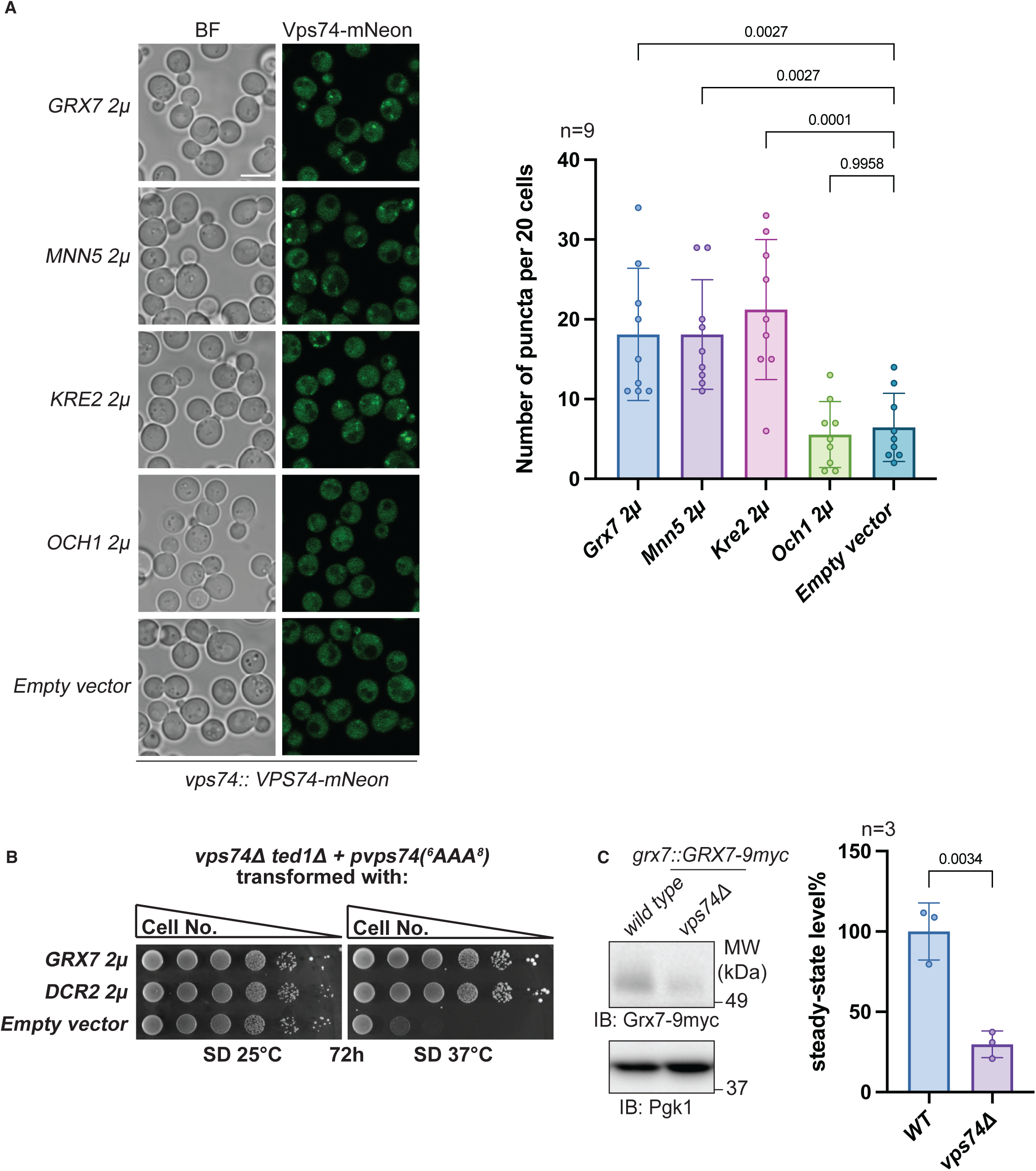
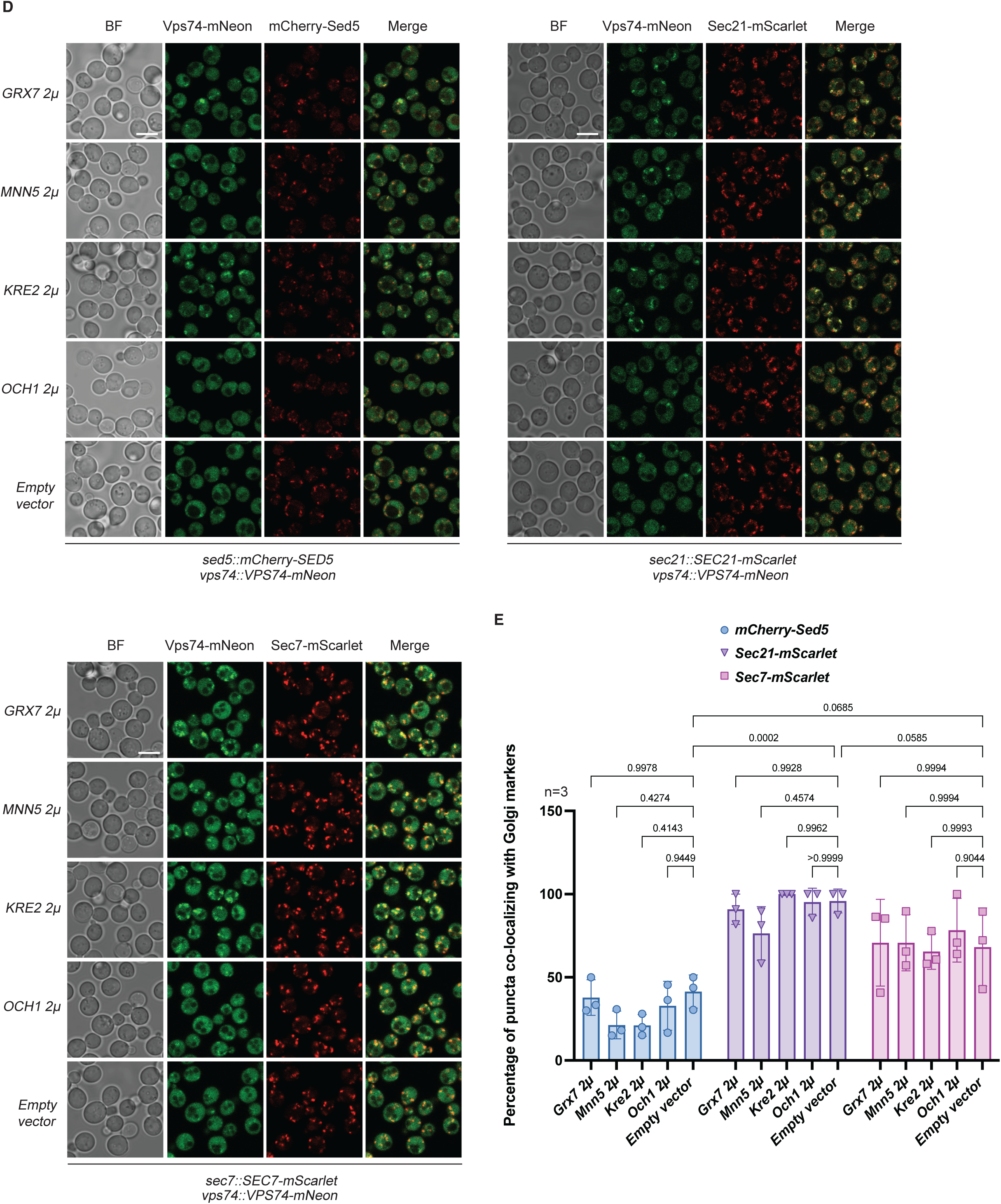
Over-expression of clients is sufficient to recruit Vps74 to Golgi membranes. (A) Cells expressing *GRX7*, *MNN5* and *KRE2* from high copy number plasmids show more Golgi puncta. Scale bar 5 μm. (B) *GRX7* is a dosage suppressor of the temperature-sensitive growth phenotype of *vps74-1 ted111* cells. 10-fold serial dilutions of the indicated yeast strains were spotted onto plates and thereafter incubated as indicated. (C) The steady-state level of Grx7p is reduced in cells lacking *VPS74*. (D) Over-expression of *GRX7*, *MNN5* and *KRE2* is sufficient to recruit Vps74-mNeon to *cis*, medial and *trans* Golgi cisternae. Scale bar 5 μm. (E) Client over-expression does not significantly alter the distribution of Vps74 across Golgi cisternae.

We recovered *GRX7* as a dosage suppressor of the temperature-sensitivity of *vps74-1 ted111* cells (Figure 2B), and like other clients, in the absence of *VPS74* the steady-state levels of Grx7 were reduced (∼20% of endogenous levels) (Figure 2C). Grx7 is a type II integral membrane protein with a short cytoplasmic N-terminal region and reportedly functions as a Golgi-localized monothiol glutaredoxin (Mesecke et al., 2008).

Possible explanations for the observed increase in puncta in cells overexpressing Grx7, Mnn5 and Kre2 include alterations to the steady-state distributions of these proteins, whereby they are found in more distal and PI4P-enriched cisternae. Alternatively, Vps74 can be recruited to the Golgi by binding directly to the N-termini of its clients. To address these scenarios, we introduced the high copy number *GRX7*, *MNN5* and *KRE2* containing plasmids into cells expressing Vps74-mNeon, and either mCherry-Sed5, Sec21-mScarlet or Sec7-mScarlet, and quantified the number of co-localizing puncta (Figure 2D). These experiments revealed that over-expression of individual clients did not alter the overall percentage of Vps74-mNeon puncta in cisternae fluorescently labelled with Sed5, Sec21 or Sec7 when compared to cells transformed with empty vector only (Figure 2D and E). To conclude, clients can play a direct role in the recruitment of Vps74 to Golgi cisternae.

### Client over-expression is not sufficient to recruit the PI4P-binding deficient and β hairpin variants of Vps74 to the Golgi

Previous studies identified a role for PI4P and the β hairpin in the recruitment of Vps74 to Golgi membranes (Wood et al., 2009). Considering the findings presented in Figure 2, we re-examined the role of these features in the recruitment of Vps74 to the Golgi by generating amino acid substitutions to the PI4P binding site at R97, W88 and K178, R181, and deleting the amino acids comprising the β hairpin (amino acids 197 – 208 (Δ197-208)) (Wood et al., 2009). The functional consequences of these variants were assessed in cells lacking *VPS74* and *TED1* (Figure 3A). Only the R97A variant of Vps74 was incapable of supporting the growth of *vps7411 ted111* cells (Figure 3A). Surprisingly, substitutions at three of the four amino acids previously reported to be important for Vps74 binding to PI4P (W88, K178 and R181) did not result in loss of function of the protein, nor were *vps7411 ted111* cells expressing these variants (W88A and K178A R181A) temperature-sensitive for growth (Figure 3B). Although *vps7411 ted111* cells expressing the β hairpin deletion variant vps74 (Δ197-208) were viable at 25°C, these cells were temperature-sensitive for growth at 37°C (Figure 3B). The β hairpin deletion variant also conferred calcofluor white sensitivity to cells (a proxy for glycosylation defects) (Ram et al., 1994), whereas *vps7411 ted111* cells expressing the W88A and K178A R181A amino acid substitution variants of Vps74 were no more sensitive to calcofluor white, than cells expressing the wildtype protein (Figure 3B). It appears that the β hairpin does not play a crucial role in the Golgi membrane recruitment of Vps74 when cells are grown at 25°C.

**Figure 3.**
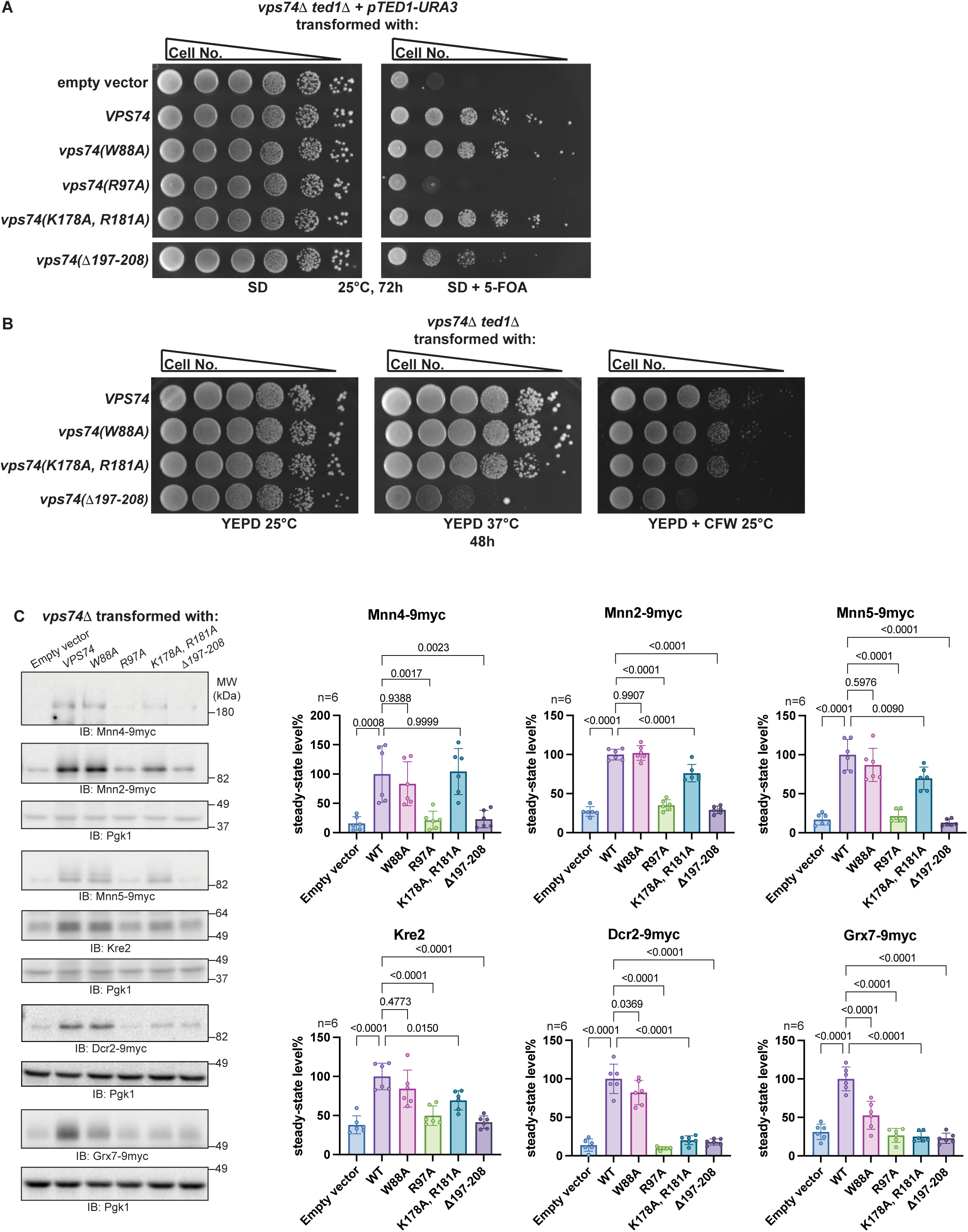
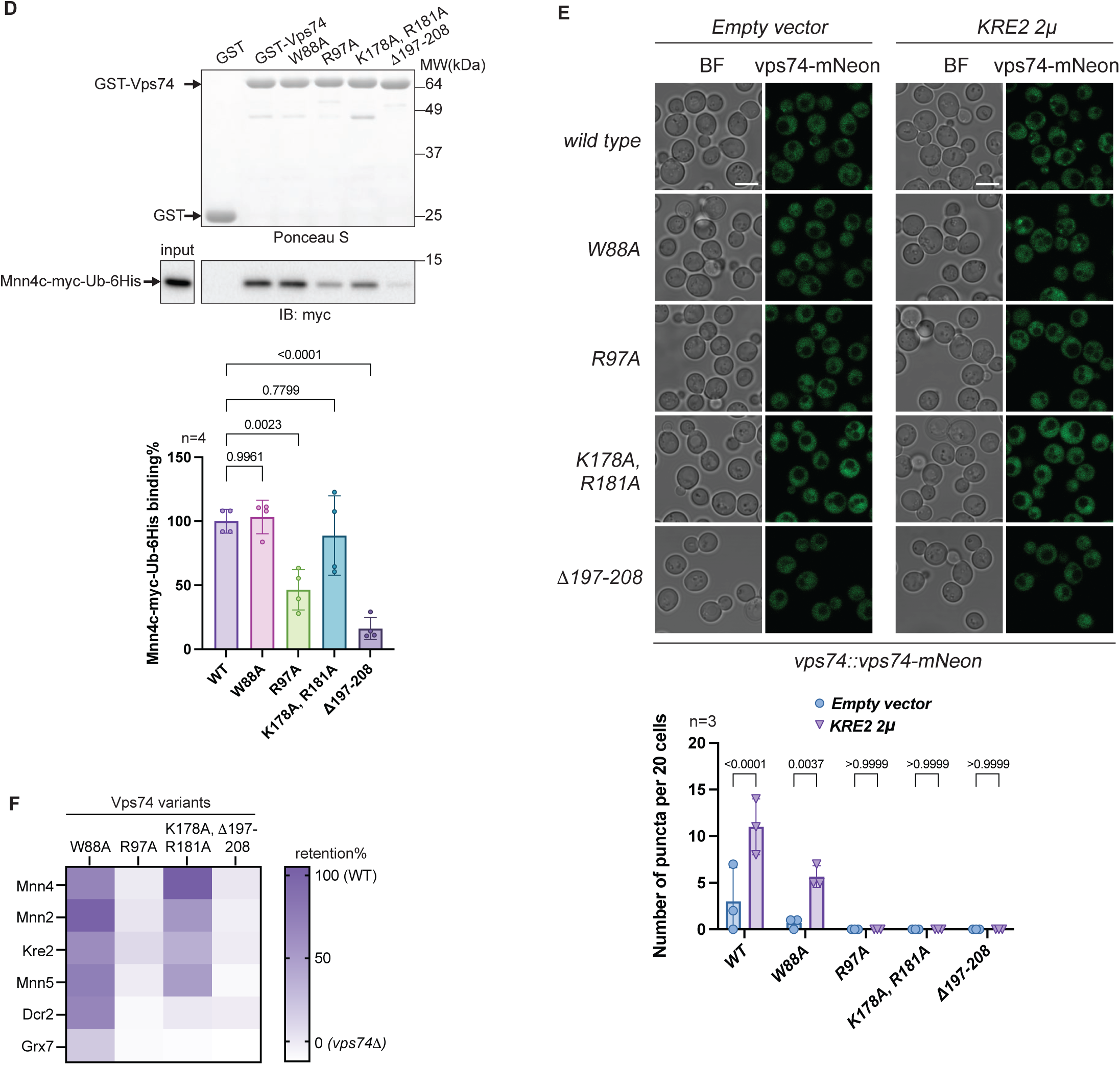
Amino acid substitutions to the PI4P-binding site and deletion of the β hairpin of Vps74 impact client-binding *in vivo* and *in vitro*. (A) The R97 variant of Vps74 cannot support the growth of *vps7411 ted111* cells. 10-fold serial dilutions of the indicated yeast strains were spotted onto plates and thereafter incubated as indicated. (B) Expression of the β hairpin deletion variant confers temperature-sensitivity for growth and calcofluor sensitivity in *vps7411 ted111* cells. 10-fold serial dilutions of the indicated yeast strains were spotted onto plates and thereafter incubated as indicated. (C) Immunoblots and quantification of the steady-state levels of clients in *vps7411* cells expressing the Vps74 variants. Pgk1 serves as a gel load control. (D) The R97, K178 R181, or β hairpin deletion (11197-208) variants show reduced binding to the N-terminus of Mnn4 in an in vitro mixing assay. (E) Over-expression of Kre2 is not sufficient to recruit the R97, K178 R181, or β hairpin deletion (11197-208) variants to Golgi membranes. Scale bar 5 μm. (F) A heat map depicting the impact of the Vps74 variants on client retention. Note that the steady-state levels of clients in cells lacking *VPS74* (Empty vector, panel C) have been subtracted.

An important caveat in the interpretation of these findings is the genetic background (*vps7411 ted111*) used to assess the function of the various amino acid variants. *TED1* encodes a remodelase that removes phosphoethanolamine from mannose 2 of glycosylphosphatidylinositol-anchored proteins (GPI-AP) in the ER, an essential function that is shared by Dcr2, which acts in the Golgi (Chen et al., 2021). Dcr2 is a Vps74 client, and thus cells lacking *VPS74* and *TED1* are not viable because Dcr2 is mislocalized and degraded in the vacuole (Chen et al., 2021). Therefore, whether a particular variant can support the growth of cells lacking *VPS74* and *TED1*, reflects the variant’s impact on the Golgi retention of Dcr2 (Chen et al., 2021). In line with this reasoning, the steady-state levels of Dcr2 were significantly reduced in the R97A, K178A R181A and β hairpin variants, comparable to those observed in cells lacking *VPS74* (Figure 3C). Presumably, the essential activity of Dcr2 can be provided by ∼ 20% of endogenous levels of the enzyme (Figure 3C), which may explain the observation that the β hairpin variant can support the growth of *vps7411 ted111* cells (Figure 3A and B). Nevertheless, these findings were recapitulated in *in vitro* mixing experiments using the Mnn4 N-terminus (Figure 3D) and revealed a direct role for the β hairpin and R97 (in the PI4P binding site) in client-binding. Thus, in addition to the reported importance of the β hairpin in Golgi membrane-binding (Cai et al., 2014) it also appears to play a key role in Vps74 client-binding, at least in the case of Mnn4.

To further interrogate the impact of the W88, R97, K178 R181 and β hairpin variants on Vps74 membrane-binding, we asked whether over-expression of Kre2 (Figure 2A and Figure EV2) was sufficient to recruit these adaptor variants to Golgi membranes (Figure 3E). However, puncta were only apparent in cells expressing wildtype *VPS74*, and to a lesser extent in cells expressing the W88A substitution variant (Figure 3E).

To explore the extent to which the Vps74 variants affected the retention of clients in the Golgi, we examined the steady-state levels of Kre2, Mnn4, Mnn2, Grx7 and Mnn5 in variant-expressing cells (Figure 3C and F). These experiments revealed that substitutions to W88 and K178, R181 had a comparatively minimal impact on the steady-state levels of clients, whereas the R97 and the β hairpin variants had the largest effect on client retention (Figure 3C and F). Although amino acid substitutions at R97, W88 and K178, R181 reportedly resulted in mislocalization of Kre2 to the vacuole (Wood et al., 2009), these variants of Vps74 (with the exception of the R97 variant, which was not tested) were not sensitive to calcofluor white (Figure 3B) unlike cells lacking the *KRE2* (Tu et al., 2008), nor did cells expressing the W88 and K187, R181 variants show a precipitous reduction in the steady-state levels of Kre2 (Figure 3C and F).

Our results on substitutions to R97, W88 and K178, R181 are difficult to reconcile with a critical role for PI4P recruiting Vps74 to Golgi membranes. Except for Grx7, the W88 variant had a comparatively modest effect on Vps74 client retention (Figure 3C and F). Similarly, except for Dcr2 and Grx7, the K178, R181 variant had a comparatively modest effect on all other clients tested (Figure 3C and F). However, as a structure of Vps74 bound to PI4P is not available, we cannot exclude the possibility that other amino acids in addition to R97 might also be critical for Vps74-PI4P binding in cells. Nevertheless, R97 and the β hairpin appear to play a role in adaptor-client-binding, as judged by *in vitro* binding studies with the N-terminus of Mnn4 (Figure 3D).

### Recruitment of Vps74 variants to Golgi membranes with a PI4P binding PH domain is not sufficient to restore client retention

To further assess whether recruitment of Vps74 to PI4P-enriched Golgi cisternae was critical for client retention, we generated a dimer of Vps74 in which the PH domain of FAPP1, which has been shown to bind to PI4P in yeast cells and to co-localize with the *trans* Golgi protein Sec7 (Faulhammer et al., 2005; Faulhammer et al., 2007), was inserted between the two monomers (Figure 4A). We opted for tethering two molecules of Vps74 as a covalent dimer is functional (Figure 1B and C). The W88A, R97A, and K178A, R181A substitutions and β hairpin deletion variant were introduced into both Vps74 monomers (Figure 4A) and were expressed from the *VPS74* locus. In cells expressing Vps74-mNeon-Vps74 and the vps74-mNeon-vps74 variants, only WT Vps74-mNeon-Vps74 and the W88A vps74-mNeon-vps74 variant showed any puncta (Figure 4B), in agreement with our observations with Vps74-mNeon and these variants (Figure 3C). By contrast, WT Vps74-mNeon-PH-Vps74 and all the vps74-mNeon-PH-vps74 variants showed puncta, consistent with the ability of the FAPP1 PH domain to bind PI4P in yeast cells (Figure 4B) (Faulhammer et al., 2005). However, when we examined the steady-state levels of two clients (Mnn5 and Kre2) in cells expressing the Vps74-mNeon-Vps74 or vps74-mNeon-PH-vps74 variants R97A, K178A, R181A and Δ197-208 (the β hairpin deletion), the vps74-mNeon-PH-vps74 variants were no better than the vps74-mNeon-vps74 variants in restoring the steady-state level of Mnn5 or Kre2 (Figure 4C). Interestingly, both the WT and W88 variant of Vps74-mNeon-PH-Vps74 were less effective in maintaining the steady-state levels of Mnn5 than their Vps74-mNeon-Vps74 counterparts, suggesting that when the adaptor was localized to more *trans* cisternae PI4P enriched cisternae, it was less efficient retaining this client in the Golgi (Figure 4C). To conclude, our findings are consistent with the hypothesis that the binding of Vps74 to PI4P is not critical for the adaptor’s function *per se*, but rather that one or more of the amino acids previously identified as being required for PI4P binding (R97 and K178, R181) are in fact part of the client-binding site. Indeed, it may be the case that directing the adaptor to more *trans* cisternae via a PH domain is detrimental, perhaps when Vps74 is occupied with PI4P the protein is less able to bind its clients.

**Figure 4.**
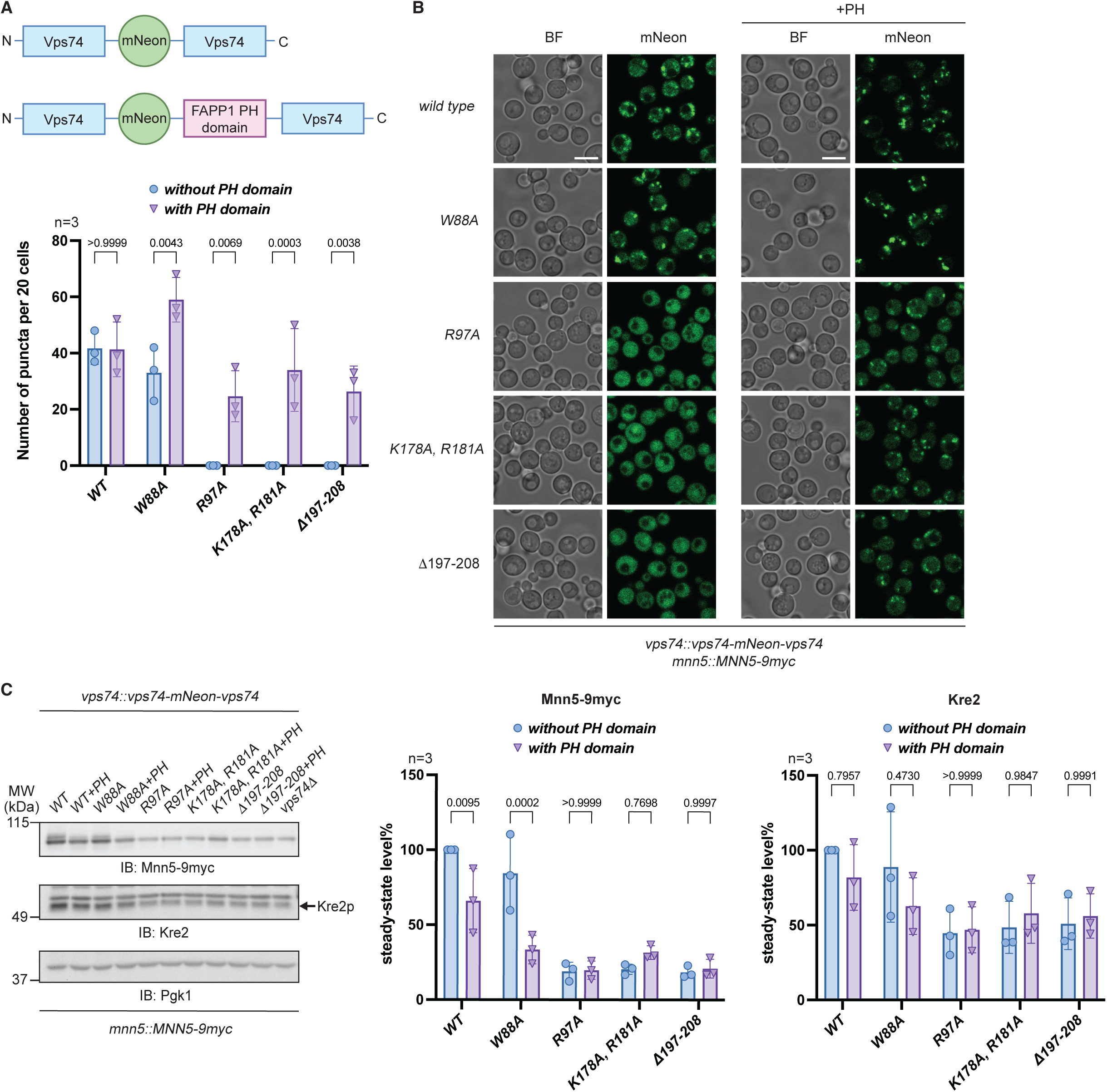
Vps74-mNeon-PH-Vps74 variants do not retain clients in the Golgi. (A) Depiction of constructs used in Panels B and C. (B) Inserting the PH domain of FAPP1 into Vps74-mNeon-Vps74 is sufficient to recruit Vps74 variants to the Golgi. Scale bar 5 μm. (C) The R97, K178 R181 and β hairpin variants of Vps74-mNeon-PH-Vps74 do not restore the steady-state levels of Mnn5 or Kre2. Pgk1 serves as a gel load control.

### The client-binding site on Vps74 includes two evolutionarily conserved adjacent unstructured loops

The prospect that the amino acid residues important for PI4P binding and the β hairpin might constitute part of the client-binding site on Vps74, prompted us to search for evolutionarily conserved, surface exposed and membrane opposing regions on Vps74 and human GOLPH3s. This analysis identified two regions on Vps74 / GOLPH3s defined by amino acids 78 – 90 and 165 – 177 (Vps74 numbering) (Figure 5A). These residues define two unstructured regions on Vps74 (hereafter referred to as Loop1 and Loop2) that flank the PI4P binding site and the β hairpin (Figure 5A and B). To identify those amino acids critical for client-binding, we used AlphaFold2. For these analyses we used the amino acid sequences of 9 client N-termini (Figure EV3A) and evaluated possible interactions of these clients with Vps74 from the AlphaFold2 predictions. This *in silico* analysis identified W88 and D90 in Loop1 and E166, W168 and Q176 in Loop2 as the most frequent presumptive contact sites between clients and Vps74 (Figure 5C).

**Figure 5.**
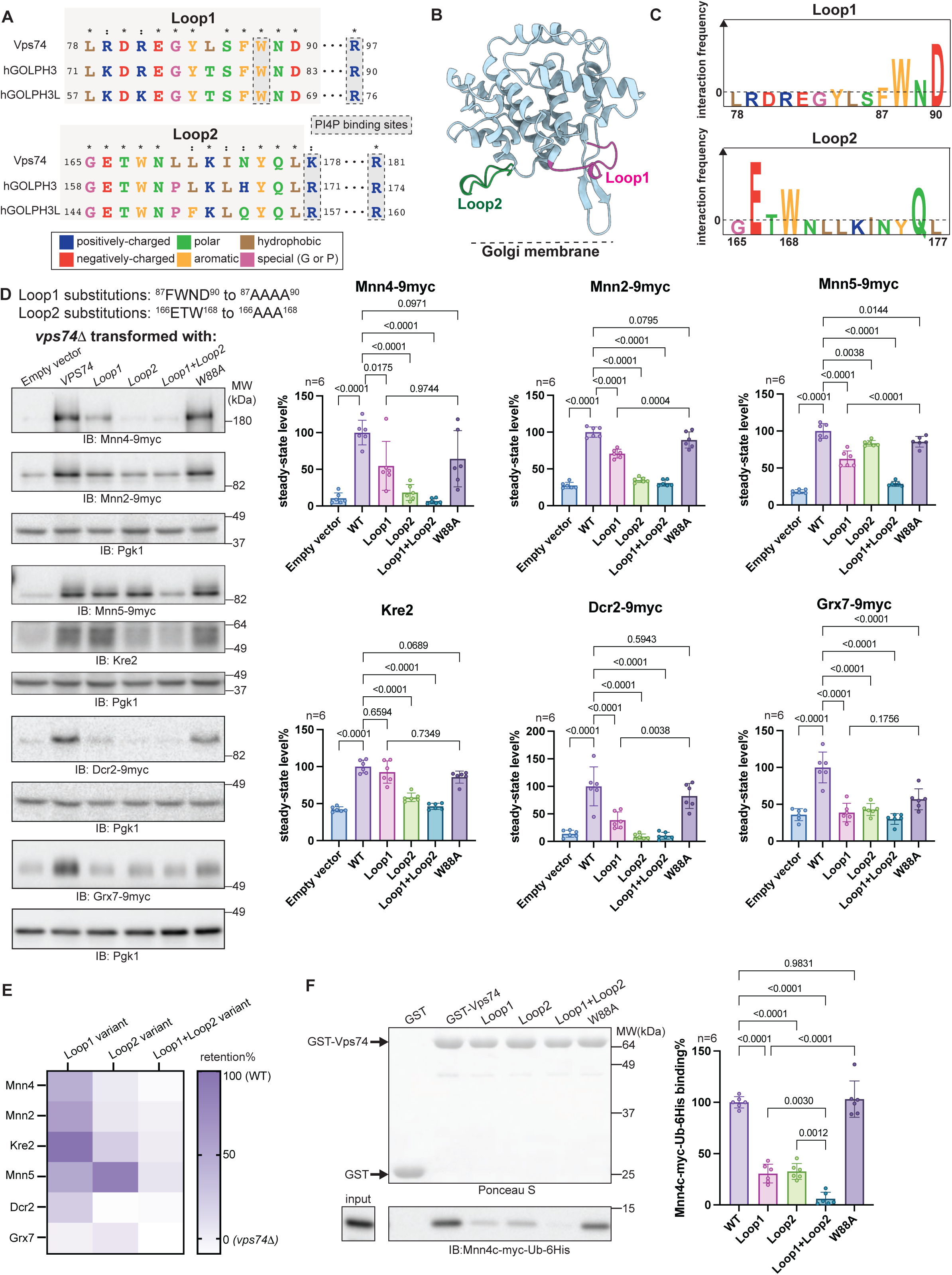
The client-binding site on Vps74 is located in two adjacent unstructured loops. (A) The GOLPH3s share two conserved, surface exposed flexible loops located on the membrane-proximal surface. Note that Loop1 and Loop2 are adjacent to the PI4P binding site. (B) A structural rendering of Vps74 in which the locations of Loop1 and Loop2 are shown relative to the Golgi membrane surface. (C) Docking client N-termini to Vps74 with AlphaFold2 identifies frequent contact sites in the client-binding adaptor interface. (D) Immunoblots and quantification of steady-state levels of clients in *vps7411* cells expressing the Loop1, Loop2, Loop1+Loop2 and W88 variants. Pgk1 serves as a gel load control. (E) A heat map summarizing the impact of the Vps74 Loop1 and Loop2 variants on client retention. Note that the steady-state levels of clients in cells lacking *VPS74* (Empty vector, panel D) have been subtracted. (F) The impact of the Loop1 and Loop2 variants on Mnn4 binding to Vps74 are recapitulated in an *in vitro* mixing assay.

**Figure 6.**
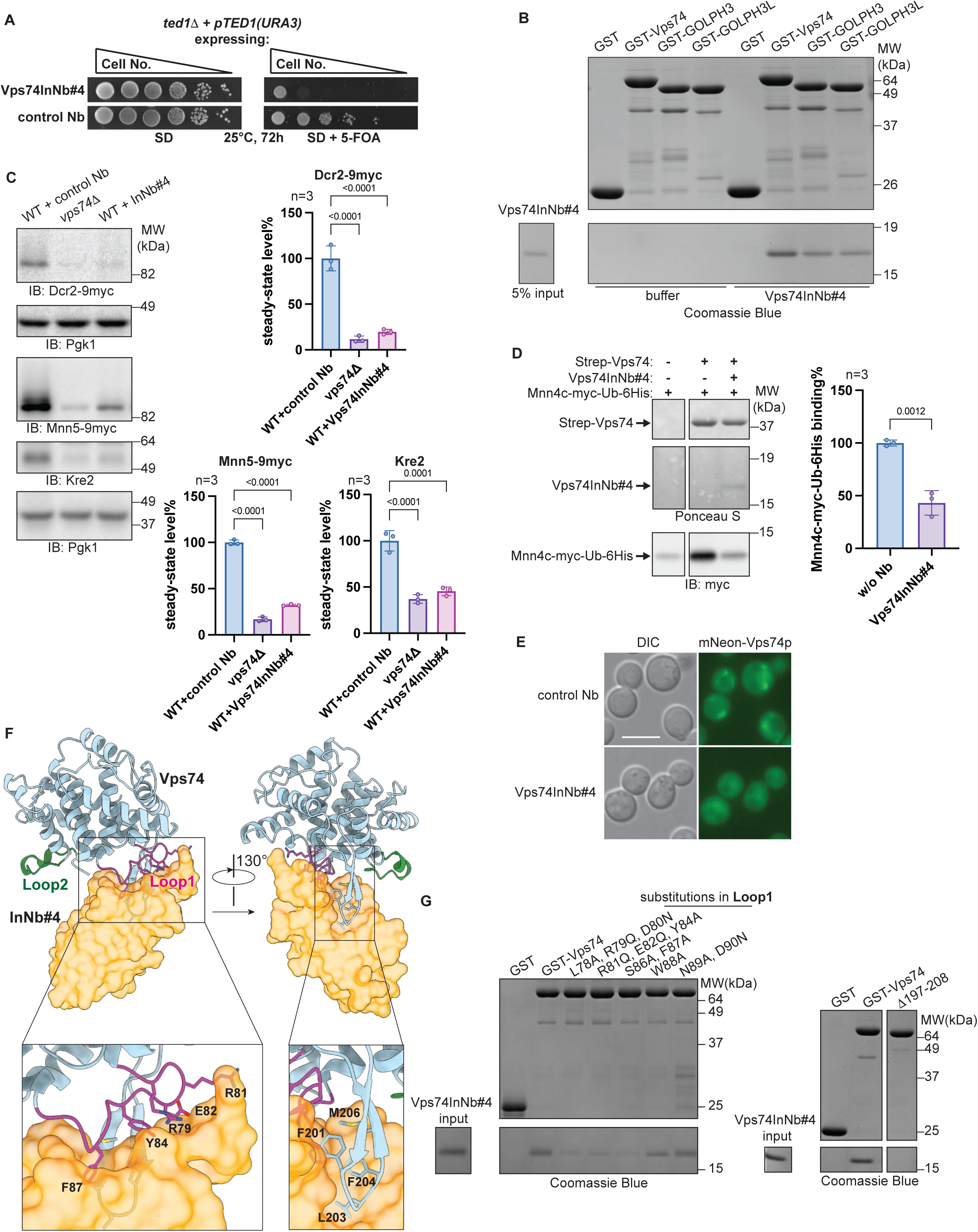
The nanobody Vps74InNb#4 corroborates identification of the client-binding site on Vps74. (A) Expression of nanobody Vps74InNb#4 inhibits the growth of cells lacking *TED1*. 10-fold serial dilutions of the indicated yeast strains were spotted onto plates and thereafter incubated as indicated. (B) Vps74InNb#4 binds to Vps74, human GOLPH3 and GOLPH3L. (C) Expression of Vps74InNb#4 in wildtype cells reduces the steady-state levels of Mnn5, Kre2 and Dcr2. Pgk1 serves as a gel load control. (D) Vps74InNb#4 reduces the binding of the N-terminus of Mnn4 to Vps74 *in vitro*. (E) Expression of Vps74InNb#4 in wildtype cells blocks Golgi membrane recruitment of Vps74. Scale bar 5 μm. (F) Vps74InNb#4 binds to Loop1 and the β hairpin of Vps74. Vps74InNb#4 was docked with Vps74 using AlphaFold3. (G) Validation of the binding interface for Vps74InNb#4 on Vps74 with site-directed variants of Vps74.

To examine the impact of substitutions in Loop1 and Loop2 on Vps74’s adaptor function, we generated three substitution variants that encompassed F87, W88, N89, D90 and E166, T167, W168 either singularly or in combination. Variant function was assessed in cells lacking *VPS74* and *TED1* and carrying a counter-selectable copy of the *TED1* gene on a plasmid (Figure EV3B and C). Of the three Vps74 amino acid substitution variants examined (^87^AAAA^90^, ^166^AAA^168^, ^87^AAAA^90^ + ^166^AAA^168^), cells expressing ^166^AAA^168^ or ^87^AAAA^90^ + ^166^AAA^168^ were unable to support the growth of *vps7411 ted111* cells (Figure EV3B). Cells expressing ^87^AAAA^90^ grew indistinguishably from wildtype and were not temperature-sensitive for growth or sensitive to calcofluor white treatment (Figure EV3C). We next expressed these three variants, as well as the W88A variant, in cells lacking *VPS74*, and examined the steady-state levels of Mnn5, Kre2, Mnn4, Mnn2, Dcr2 and Grx7 by immunoblotting (Figure 5D). Quantification of these data revealed that the combined substitutions to Loop1 and Loop2 (^87^AAAA^90^ + ^166^AAA^168^) had the most significant impact on the steady-state levels of all clients tested, while W88A was the least impactful (Figure 3C and F, and Figure 5D and E). The identification of W88 as a presumptive client-binding residue is puzzling but likely reflects the relative robustness of our *in-silico* method. Based on the experiments conducted herein, W88 is unlikely to be imperative for PI4P- or client-binding.

In agreement with our genetic assessment, in cells expressing the loss of function Loop2 variant ^166^AAA^168^ (Figure EV3B), the steady-state level of Dcr2 was substantially reduced (Figure 5D), whereas for the other clients examined the steady-state levels of Mnn4 and Mnn2 were affected to the greatest extent by these amino acid substitutions (Figure 5D and E). In contrast, the steady-state levels of Mnn5 were largely unaffected by amino acid substitutions to Loop2 whereas Kre2 retention was most significantly impacted by substitutions to Loop2 (Figure 5D and E). The effect of the combined Loop1+Loop2 substitutions appears to be additive, as cells expressing this variant showed a greater reduction in client steady-state levels than cells expressing either the Loop1 or Loop2 variants singularly (Figure 5D and E). The findings on Mnn4 steady-state levels in the various Vps74 variants could be replicated in an *in vitro* binding assay, using bacterially expressed Vps74 and the N-terminus of Mnn4 (Figure 5F).

In sum, these studies define the client-binding site on Vps74 and reveal that not all clients are similarly affected by amino acid substitutions at Loop1 and Loop2 (Figure 5E). The variable effect of Loop1 and Loop2 amino acid substitutions on client retention may explain in part how Vps74 (and other GOLPH3s) can accommodate client N-termini of differing amino acid composition and lengths (Figure EV3A) (Rizzo et al., 2021; Tu et al., 2008; Wang et al., 2020; Welch et al., 2021).

### A nanobody against Vps74 corroborates identification of the client-binding site

In an orthogonal approach to our use of amino acid substitutions to define the client-binding site, we isolated a suite of Vps74-binding nanobodies (McMahon et al., 2018). Plasmids expressing N-terminally ubiquitin-tagged nanobodies were introduced into yeast cells carrying a deletion of *TED1* and a counter-selectable copy of *TED1* linked to the *URA3* gene (Figure EV4A and B). Fusion to ubiquitin was necessary to increase the expression of nanobodies in the cytoplasm, however endogenous cytoplasmic deubiquitinases cleaved ubiquitin, preserving the N-terminal sequence of the nanobody (Figure EV4C). In this assay, cells expressing a nanobody that blocked an essential property on Vps74 would not be able to grow in *ted111* cells as neutralization of Vps74 activity would phenocopy *ted111 vps7411* cells (Figure EV4B). We chose one such inhibitory nanobody termed Vps74InNb#4 for further characterization (Figure 6A), as it also bound to human GOLPH3 and GOLPH3L, reasoning that it might bind to an evolutionarily conserved and therefore functionally significant region on the GOLPH3s (Figure 6B).

In agreement with its dominant negative effect in *ted111* cells, when Vps74InNb#4 was expressed in wildtype cells, the steady-state levels of clients were significantly reduced as judged by immunoblots for Dcr2, Mnn5 and Kre2 (Figure 6C). The reduction of steady-state levels of Dcr2, Mnn5 and Kre2 was a direct result of Vps74InNb#4 binding to Vps74 (Figure EV4C). Vps74InNb#4 also reduced the association of Mnn4 with Vps74 in an *in vitro* mixing assay (Figure 6D). Thus, Vps74InNb#4 bound to a membrane opposing surface on Vps74, a property that also blocked the binding of Vps74 to Golgi membranes (Figure 6E) and resulted in the mislocalization of clients to the vacuole (Figure EV4D).

To validate that Vps74InNb#4 occupied the client-binding site, we docked the nanobody with Vps74 using AlphaFold3 (Figure 6F) and verified the interaction interface on Vps74 using site-directed variants in an *in vitro* mixing assay (Figure 6G and Figure EV4E). To conclude, Vps74InNb#4 bound to the β hairpin and to Loop1, but not to Loop2. Nevertheless, the interaction of Vps74InNb#4 with Vps74 was sufficient to prevent the adaptor from binding to Golgi membranes, and directly to Mnn4 in *in vitro* mixing experiments (Figure 6D and E). These data corroborate our interrogation of Loop1 and Loop2 by site directed mutagenesis (Figure 5).

### The Arf1-GTP and Sac1 binding sites overlap with and exclude client-binding to Vps74

Vps74 has been shown to bind to the PI4P phosphatase Sac1, whereupon Vps74 reportedly stimulates the enzyme’s activity (Cai et al., 2014; Wood et al., 2012). Critically, the binding site on Vps74 for Sac1 encompasses Loop2 (Cai et al., 2014) coinciding with part of the client-binding site we identify in this study (Figure 5). We have previously shown that Vps74 binds to the small GTPase Arf1 in its GTP-bound state (Tu et al., 2012). Unlike the Vps74 – Sac1 association, the location of the Arf1-GTP binding site on Vps74 and the functional significance of this interaction has remained elusive. Nonetheless, Vps74 / GOLPH3 binds to the Golgi vesicle coat complex coatomer (Eckert et al., 2014; Tu et al., 2012) and Arf1-GTP is critical for COPI vesicle formation (Orci et al., 1993). These findings are consistent with a role for Arf-GTP in the incorporation of Vps74 and its clients into COPI coated vesicles.

We took advantage of the availability of Vps74InNb#4 to investigate the relationship between Vps74, and its binding partners - Sac1 and Arf1-GTP. In *in vitro* mixing experiments Sac1 reduced Mnn4 binding to Vps74 by ∼ 85% (Figure 7A). These results concur with the data presented in Figure EV5A, where we show that substitutions in Loop2, but not Loop1, reduce the binding of Sac1 to Vps74 in an *in vitro* mixing assay. These findings were corroborated with Vps74InNb#4, which preferentially binds to the β hairpin and Loop1 residues (Figure 6F and G, and Figure 7B). Thus, when Vps74 is bound to Sac1, the adaptors capacity to bind its clients is reduced.

**Figure 7.**
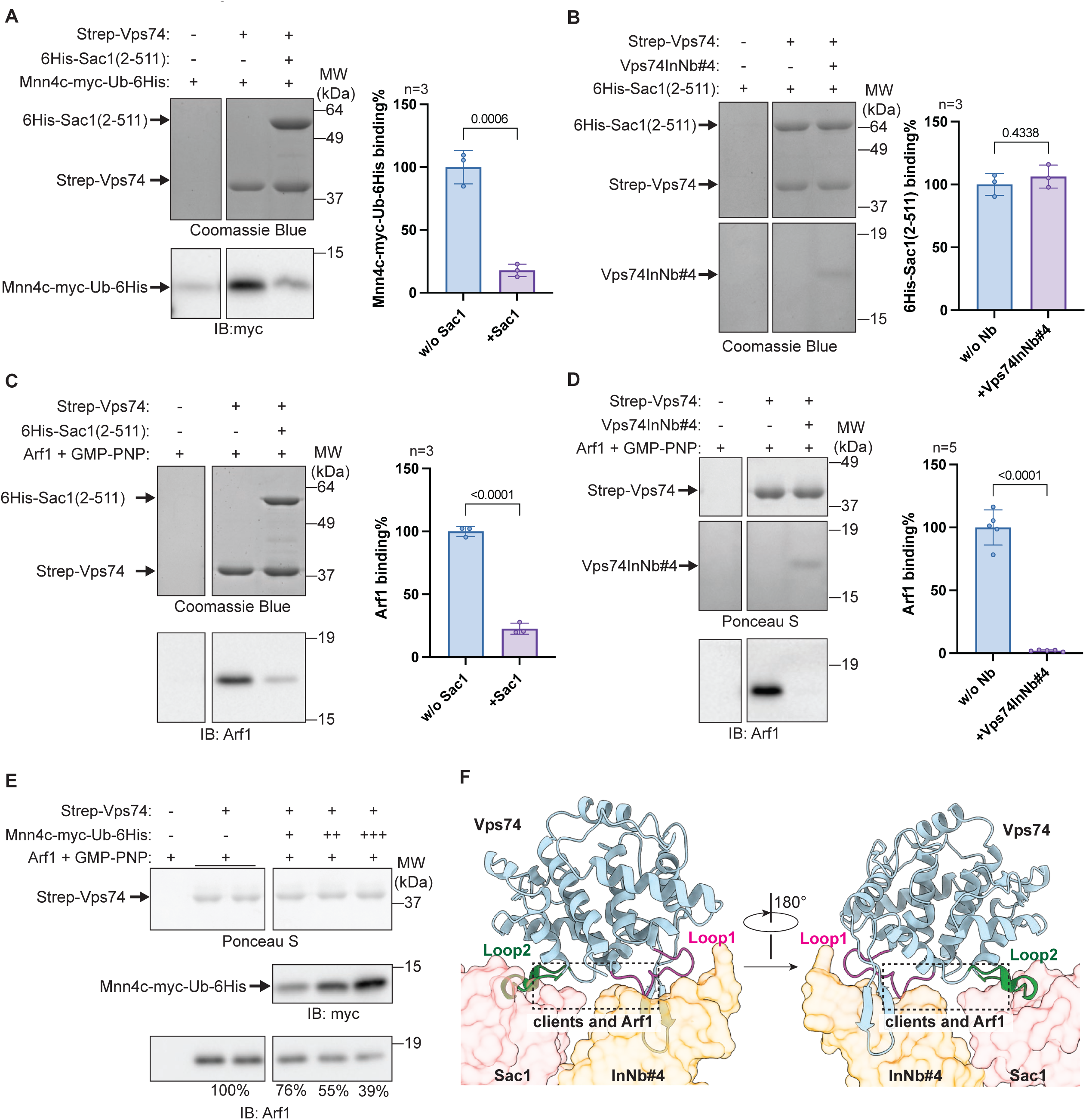
Client, Arf1 and Sac1 binding sites overlap on Vps74. (A) Sac1 competes with Mnn4 for binding to Vps74. (B) Vps74InNb#4 does not block Sac1 binding to Vps74. (C) Sac1 competes with Arf1-GTP for binding to Vps74. (D) Vps74InNb#4 blocks binding of Arf1-GTP to Vps74. (E) Arf1-GTP competes with Mnn4 for binding to Vps74. (F) The location of the client, Sac1 and Arf1-GTP binding sites on Vps74.

Interestingly, the binding site on Vps74 for Arf1-GTP and Sac1 also overlap. Sac1 binding to Vps74 resulted in a ∼ 80% reduction in the binding of Arf1-GTP (Figure 7C). Overlapping binding sites for Sac1 and Arf1-GTP were validated using site directed mutants of Vps74 in *in vitro* mixing experiments (Figure EV5A and B). In contrast to Sac1, Vps74InNb#4 blocked Arf1-GTP binding to Vps74 (Figure 7D). This finding agrees with the AlphaFold3 predicted structure of Vps74InNb#4 bound to Vps74, which revealed that the nanobody binds to Loop1 and the β hairpin (Figure 6F and G), and with the *in vitro* mixing experiments where increasing amounts of the Mnn4 N-terminus reduced Arf1-GTP binding to Vps74 (Figure 7E).

To conclude, the data presented in Figure 7 and Figure EV5 indicate that the binding sites for clients, Sac1 and Arf1 overlap with one another (Figure 7F). Whilst we have not measured the K_D_ for Sac1, Arf1-GTP or clients on Vps74, the *in vitro* mixing data we present here is consistent with Sac1 binding more robustly to Vps74 in cells than either Arf1-GTP or client N-termini. However, the extent to which Sac1 interferes with client and Arf1-GTP binding to Vps74 may be negligible under normal growth conditions, as Sac1 is predominantly an ER resident (Faulhammer et al., 2005; Faulhammer et al., 2007), whereas Vps74 is found on the Golgi membranes and in the cytoplasm – as is also the case of Arf1. Reportedly, Sac1 is found in the Golgi in cells deprived of glucose, rather than the ER (Faulhammer et al., 2005). It is therefore possible that interactions between Sac1 and Vps74 are primarily restricted to changes in the cell’s growth parameters, such as when environmental glucose levels become limiting. The finding that Arf1-GTP and client-binding are exclusive of one another is difficult to reconcile with a role for Arf1-GFP in facilitating client selection by Vps74. Rather, Arf1-GTP may function to prevent client-free adaptors from being incorporated into nascent COPI vesicles upon hydrolysis of GTP.

## DISCUSSION

We report the identification of a single membrane opposing surface that mediates the binding of clients, Sac1 and Arf1-GTP to the Golgi adaptor protein yGOLPH3 - Vps74. This binding site is comprised of evolutionarily conserved residues that encompass two unstructured regions (designated Loop1 and Loop2), the PI4P binding site, and the β hairpin. These findings were corroborated using a nanobody (Vps74InNb#4) that binds to Loop1 and the β-hairpin of Vps74 in *in vitro* binding studies using the Mnn4 N-terminus, Sac1 and Arf1-GTP. As such, we anticipate that the client-binding site we identify on Vps74 will likely play an equivalent role in GOLPH3 family proteins.

We examined the steady-state levels of six Vps74 clients, of which four are glycosyltransferases, one a monothiol glutaredoxin (Grx7) and one a GPI-AP remodelase (Dcr2). These client N-termini are varied in the amino acid sequence composition, length and cisternal distribution. Our study reveals that clients can have differing requirements on the Vps74 binding interface. For example, the Golgi retention of Mnn4, Mnn2 and Dcr2 are particularly sensitive to amino acid substitutions in Loop2. While Mnn5 retention is particularly sensitive to amino acid substitutions in Loop1+Loop2 whereas Kre2 retention is largely unaffected by amino acid substitutions in Loop1 and only moderately affected by substitutions to Loop2 or Loop1+Loop2 (Figure 5). These findings may help to reconcile how the GOLPH3 adaptor proteins can accommodate client N-termini of varying lengths and amino acid compositions (Rizzo et al., 2021; Schmitz et al., 2008; Tu et al., 2008; Welch et al., 2021).

Our findings on the requirement for amino acids previously shown to be important for PI4P- and membrane-binding (via the β hairpin) are more difficult to interpret as client-binding can also contribute to Vps74 membrane-binding (Figure 2A and D). Thus, loss of membrane association *in vivo* (by fluorescence microscopy) can no longer be used as a direct indication of a loss of PI4P-binding, or a loss of membrane binding via the β hairpin. Much of our data were obtained by examining the steady-state levels clients, it is therefore not possible to uncouple the effects of loss of Vps74 membrane association from direct loss of client-binding. Support for the PI4P-binding site in client-binding comes from examining the steady-state levels of clients in Vps74 K178, R181 variant expressing cells. Whilst the K178, R181 variant had little impact on the steady-state levels of the glycosyltransferases examined (*in vivo* or *in vitro*), this variant did significantly reduce the steady-state levels of Dcr2 and Grx7 (Figure 3C and F) despite the amino acid sequences of their N-termini being quite divergent (Figure EV3A). In line with the requirement of R97 to support the growth of *vps7411 ted111* cells (Figure 3A and B), client retention was universally affected by a substitution with Ala at this position (Figure 3C and F). This would be expected if this residue was critical for membrane-binding via PI4P. However, *in vitro* mixing experiments with Mnn4, revealed that R97 was important for Vps74-binding to this client (Figure 3D). Similarly, a critical role for the β hairpin in the membrane association of Vps74 is also unclear. Deletion of the β hairpin did not result in loss of function of Vps74 (Figure 3A), although cells expressing this variant were temperature-sensitive and calcofluor white sensitive (Figure 3B). Nevertheless, the β hairpin deletion variant of Vps74, like the R97 variant, had the greatest impact on the steady-state levels of all clients examined (Figure 3C and F). In addition, in *in vitro* mixing experiments with the Mnn4 N-terminus, recombinant Vps74 lacking the β hairpin was significantly compromised in its capacity to bind directly to this client (Figure 3D).

To conclude, our findings with the PI4P and β hairpin variants are not consistent with a crucial or exclusive role for these regions of Vps74 in Golgi membrane-binding. Rather, it may be the case that they play a dual role – that includes client-binding and membrane-binding. The data we present herein do not allow us to distinguish between these two possibilities. Nor was it possible for us to ascertain whether PI4P-binding is prerequisite or co-requisite for adaptor-client-binding. In addition, our amino acid substitution variants and *in silico* analysis did not allow us to distinguish the orientation of client N-termini on Vps74 (i.e., whether the N-terminus of a client is orientated towards Loop1 and the β hairpin, or towards Loop2). Lastly, the inhibitory nanobody Vps74InNb#4 corroborated our identification of the Mnn4-binding site on Vps74 (Figure 6D and F).

Our application of Vps74InNb#4 in *in vitro* mixing assays with Sac1, Arf1-GTP and the Mnn4 N-terminus was instrumental in establishing that Mnn4 competes for binding to Vps74 with Sac1 and Arf1-GTP. This may also be the case for other clients more generally, but this will require the use of human GOLPH3s and their clients (for example), as these interactions appear to be more robust than between their yeast counterparts (Rizzo et al., 2021; Schmitz et al., 2008; Tu et al., 2008; Welch et al., 2021). Moreover, Sac1 and Arf1-GTP also compete with one another for binding to Vps74 suggesting that in cells these proteins may function in part to negatively regulate client / adaptor interactions. These findings were entirely unexpected. What then is the physiological significance of the binding of Arf1-GTP to Vps74? Our biochemical studies revealed that the binding-site for Arf1-GTP and clients overlap and compete with one another for binding to Vps74 (Figure 7E). Perhaps the association of Arf1-GTP with Vps74 serves to prevent futile incorporation of client-less adaptor into COPI coated vesicles as hydrolysis of GTP would release any Arf1-bound Vps74 from Golgi membranes.

## EXPANDED VIEW FIGURE LEGENDS

**Figure EV1.**
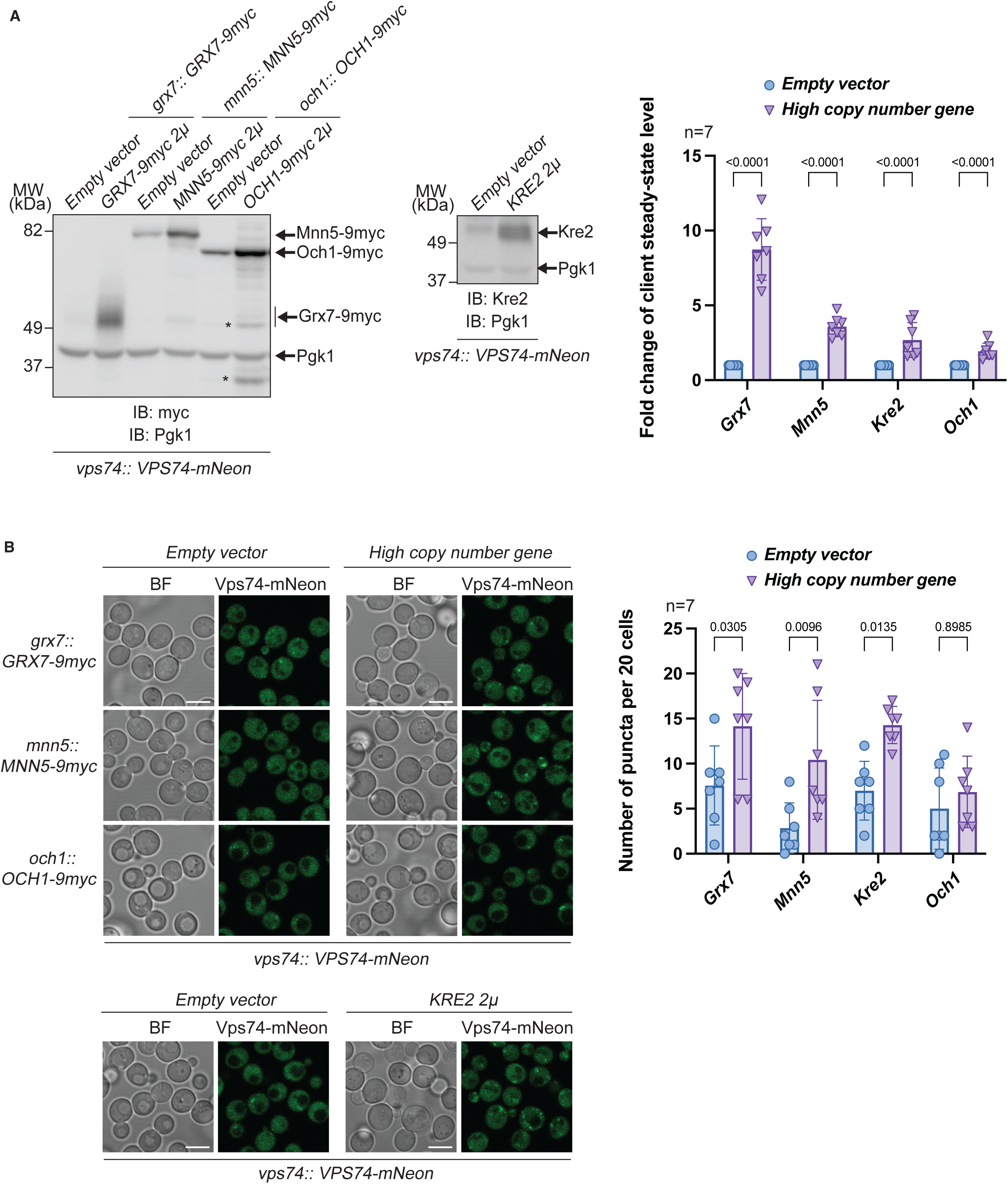
Over-expression of epitope-tagged clients increases Vps74’s binding to Golgi membranes. (A) Immunoblot and quantification of steady-state levels of clients from the indicated yeast strain. Pgk1 serves as a gel loading control. Note that Och1 is not a Vps74 client. The * indicate major degradation products of Och1 upon over-expression. (B) Cells carrying high copy number plasmids encoding *KRE2*, *GRX7-9myc* and *MNN5-9myc* have increased Vps74-mNeon binding to Golgi membranes.

**Figure EV2.**
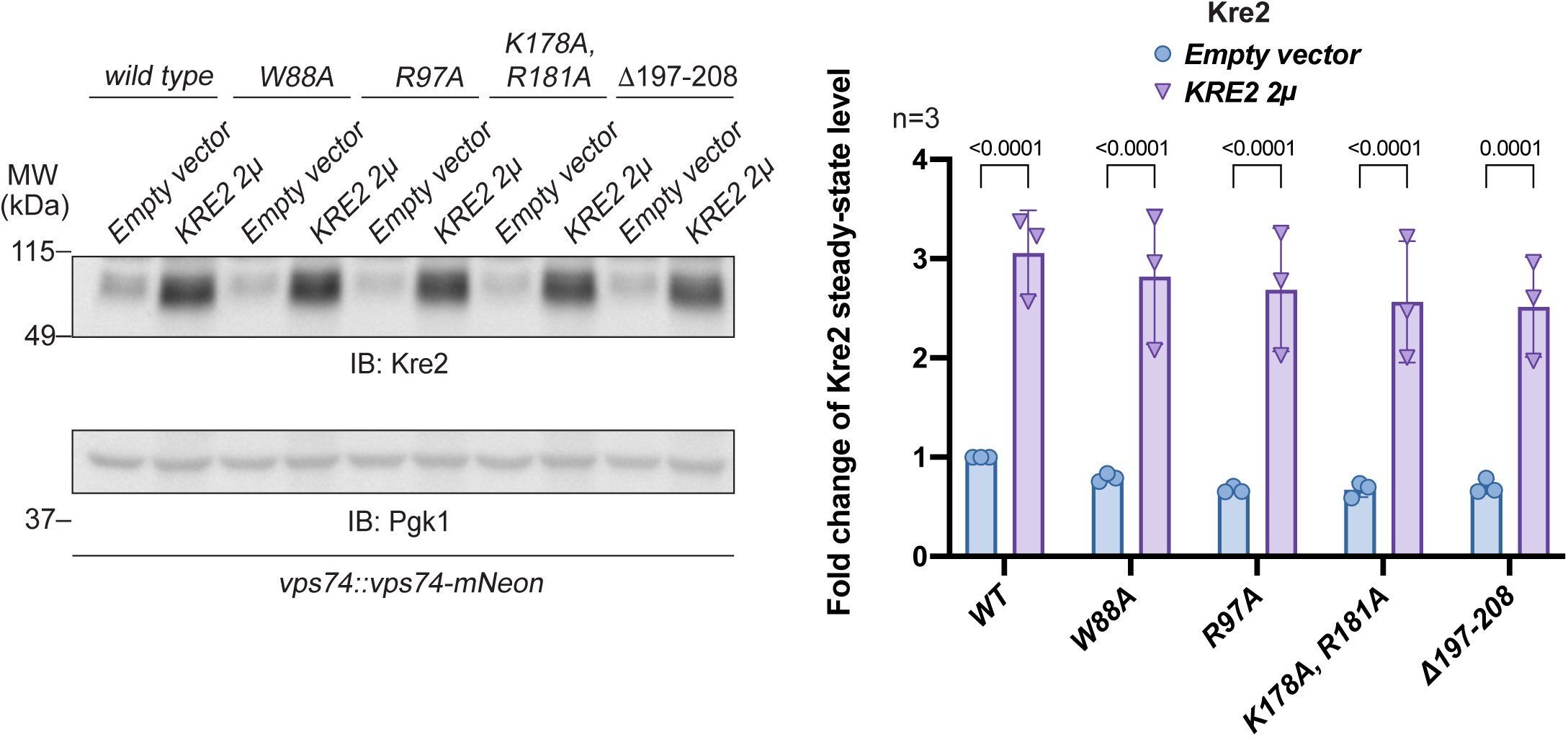
The steady-state levels and quantification of Kre2 in the W88, R97, K178 R181 and β hairpin variants of Vps74-mNeon. Pgk1 serves as a gel load control.

**Figure EV3.**
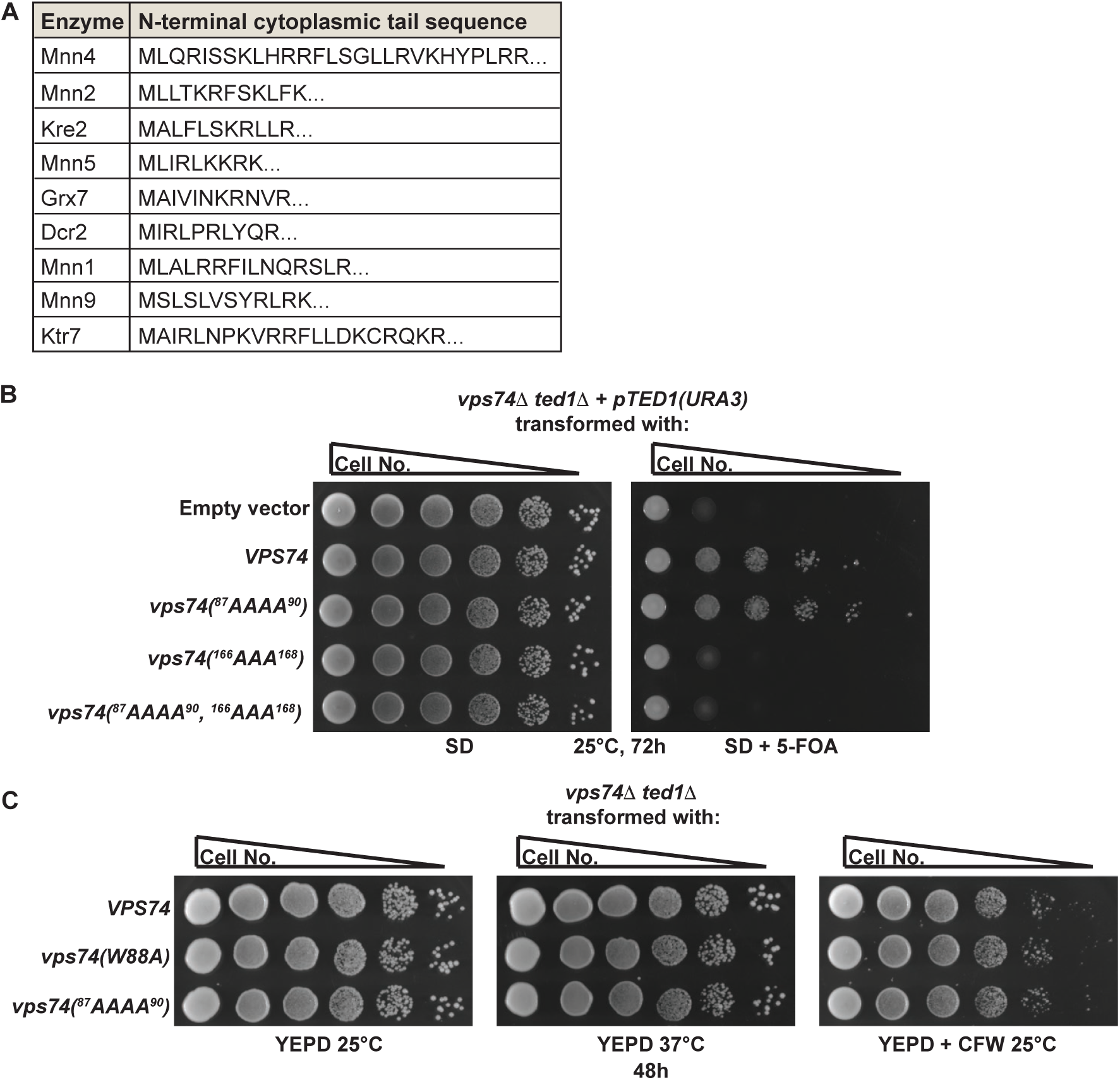
Functional analysis of amino acid substitutions to the client-binding site on Vps74. (A) Amino acid sequence alignment of the N-termini for the nine clients used in the AlphaFold2 docking analysis with Vps74. (B) Amino acid substitutions in Loop2 and Loop1+2 cannot support the growth of *vps7411 ted111* cells. 10-fold serial dilutions of the indicated strains were spotted onto plates and incubated as indicated. (C) The W88 and Loop1 variants of Vps74 are neither temperature-sensitive for growth nor sensitive to calcofluor (CFW) treatment. 10-fold serial dilutions of the indicated strains were spotted onto plates and incubated as indicated.

**Figure EV4.**
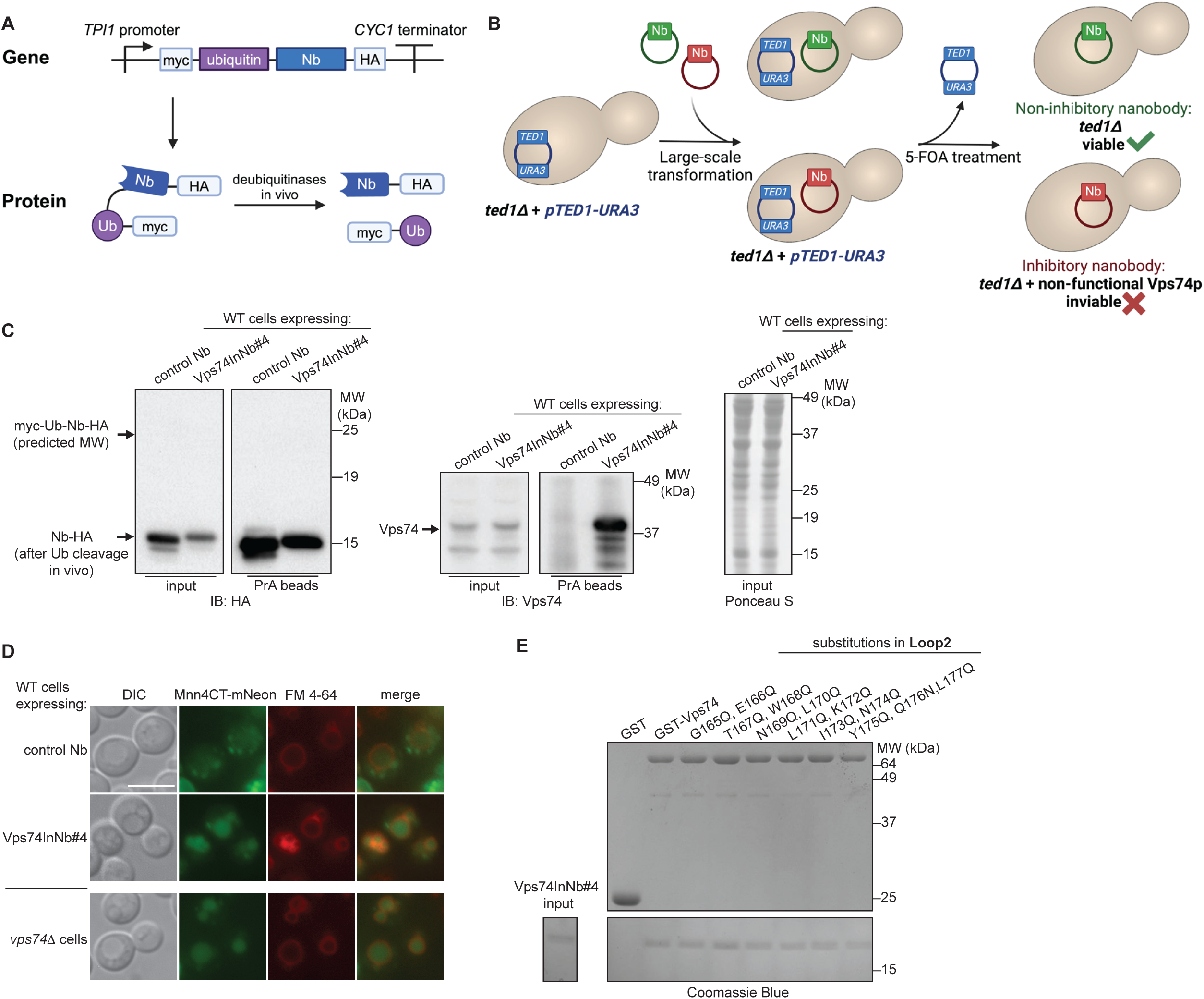
Identification and characterization of nanobodies that block the function of Vps74. (A) Scheme for the expression of nanobodies in the cytoplasm of yeast cells. (B) Scheme for the selection of plasmids expressing nanobodies that block the function of Vps74 in cells. (C) Vps74InNb#4 identified from (B) is expressed in yeast cells, cleaved to release ubiquitin, binds to Protein A (PrA) beads and can co-immune precipitate Vps74 from whole cell extracts. (D) Expression of Vps74InNb#4 in wildtype yeast cells causes mislocalization of the client reporter protein Mnn4CT-mNeon to the vacuole. CT denotes the N-terminal cytoplasmic tail and transmembrane domain, respectively. Scale bar 5 μm. (E) Amino acid substitutions in Loop2 do not affect binding of Vps74InNb#4 to Vps74.

**Figure EV5.**
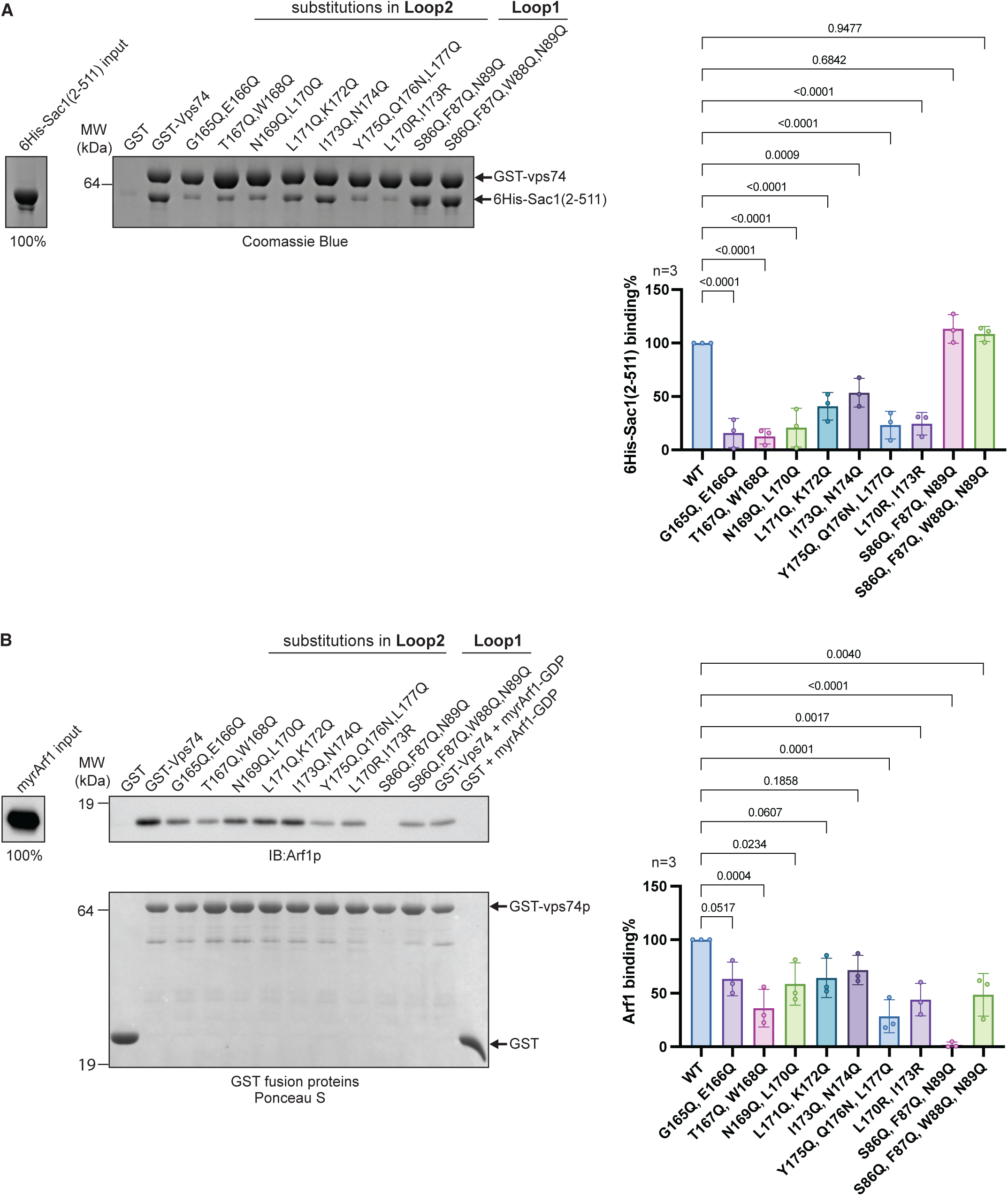
Amino acid substitutions in Loop1 and Loop2 define the binding sites for Sac1 and Arf1-GTP on Vps74. (A) *In vitro* mixing experiments using bacterially expressed purified proteins of the indicated variants with the bacterially expressed purified soluble domain of Sac1. Quantification from binding experiments is shown to the right of the binding data. (B) *In vitro* mixing experiments using bacterially expressed purified proteins of the indicated variants with bacterially expressed purified N-myristoylated Arf1-GTP. Quantification from binding experiments is shown to the right of the binding data.

